# Sequence analysis and structural predictions of lipid transfer bridges in the repeating beta groove (RBG) superfamily reveals past and present domain variations affecting form, function and interactions of VPS13, ATG2, SHIP164, Hobbit and Tweek

**DOI:** 10.1101/2022.07.06.498983

**Authors:** Tim P Levine

## Abstract

Lipid transfer between organelles requires proteins that shield the hydrophobic portions of lipids as they cross the cytoplasm. In the last decade a new structural form of lipid transfer protein (LTP) has been found: long hydrophobic grooves made of beta-sheet that bridge between organelles at membrane contact sites. Eukaryotes have five families of bridge-like LTPs: VPS13, ATG2, SHIP164, Hobbit and Tweek. These are unified into a single superfamily through their bridges being composed of just one domain, called the repeating beta groove (RBG) domain, which builds into rod shaped multimers with a hydrophobic-lined groove and hydrophilic exterior. Here, sequences and predicted structures of the RBG superfamily were analyzed in depth. Phylogenetics showed that the last eukaryotic common ancestor contained all five RBG proteins, with duplicate VPS13s. These appear to have arisen in even earlier ancestors from shorter forms with 4 RBG domains. The extreme ends of most RBG proteins have amphipathic helices that might be an adaptation for direct or indirect bilayer interaction, although this has yet to be tested. The one exception to this is the C-terminus of SHIP164, which instead has a coiled-coil. Finally, almost the entire length of the exterior surfaces of the RBG bridges are shown to have conserved residues, indicating sites for partner interactions almost all of which are unknown. These findings can inform future cell biological and biochemical experiments.

## Introduction

In the last two decades there has been a transformation in our understanding of how membrane-bound organelles of eukaryotic cells interact with each other. Text books tend to emphasize the linear pathways of secretion and endocytosis, considering organelles apart from each other and those that do not participate in vesicular traffic (mitochondria, lipid droplets, peroxisomes, plastids). This picture has increasingly been falsified by finding individual proteins that bind to two organelles at the same time, bridging the cytoplasmic gaps between them. A major activity that is found where two organelles interact is the transfer of lipids (Prinz *et al*., 2020), which can be independent of vesicular traffic (Pagano, 1990; Baumann *et al*., 2005). Lipid transport between membranes involved lipid transfer proteins (LTPs). The first discovered LTPs all have globular domains with an internal pocket specialized to shield one lipid (or possibly two) at a time (Chiapparino *et al*., 2016). Such domains can be anchored at a membrane contact site, and then shuttle back-and-forth between donor and acceptor membranes to transfer or exchange selected cargoes (Egea, 2021).

Subsequently, LTPs with an elongated rod-like structure ∼20 nm long were found in bacteria, with a “U”-shaped cross-section, the internal surface of which is entirely hydrophobic while the external surface is hydrophilic (Takeda *et al*., 2003; Suits *et al*., 2008). This allows lipids to slide between compartments along relatively static bridges (Sherman *et al*., 2018). Cytoplasmic bridge-like LTPs with a similar hydrophobic groove were then discovered in eukaryotes: VPS13 and ATG2 are distantly related proteins that form rods approximately 20 and 15 nm long containing 3000-4000 aa and 1500-2000 aa respectively (Figure 1A) (Kumar *et al*., 2018; Osawa and Noda, 2019; Valverde *et al*., 2019; Ugur *et al*., 2020; Dziurdzik and Conibear, 2021; Leonzino *et al*., 2021). VPS13 and ATG2 transfer phospholipid rapidly *in vitro* (von Bülow and Hummer, 2020; Zhang *et al*., 2022), and they are required for rapid growth of the yeast prospore membrane and autophagosomes (respectively), both of which have few embedded proteins, indicating delivery of lipid in bulk (Lees and Reinisch, 2020). A pivotal observation is that VPS13 function is inhibited by converting the lining of one segment of the rod from hydrophobic to charged residues (Li *et al*., 2020) (Figure 1A). This indicates that lipids must pass every point of the tube, strongly supporting the bridge model where lipids flow along VPS13 and by implication any related LTP bridge. Three further eukaryotic bridge-like LTPs have since been identified on the basis of distant sequence homology to VPS13/ATG2 using HHpred (Soding, 2005), and supported by structural predictions by AlphaFold (Jumper *et al*., 2021). They are: SHIP164 (name in humans also called UHRF1BP1L, with a close human homolog UHRF1BP, and a plant homolog: amino-terminal region of chorein); Tweek (name from *Drosophila*, in human: BLTP1 (newly named BLTP1 by the Human Genome Gene Nomenclature Committee, see https://www.genenames.org, previously KIAA1109 or FSA, in yeast: Csf1); and Hobbit (name from *Drosophila*; in human: BLTP2 (new name as above), previously KIAA0100, in yeast: Fmp27/Ypr117w newly named Hob1/2, in plants: SABRE, KIP and APT1).

**Figure 1:**
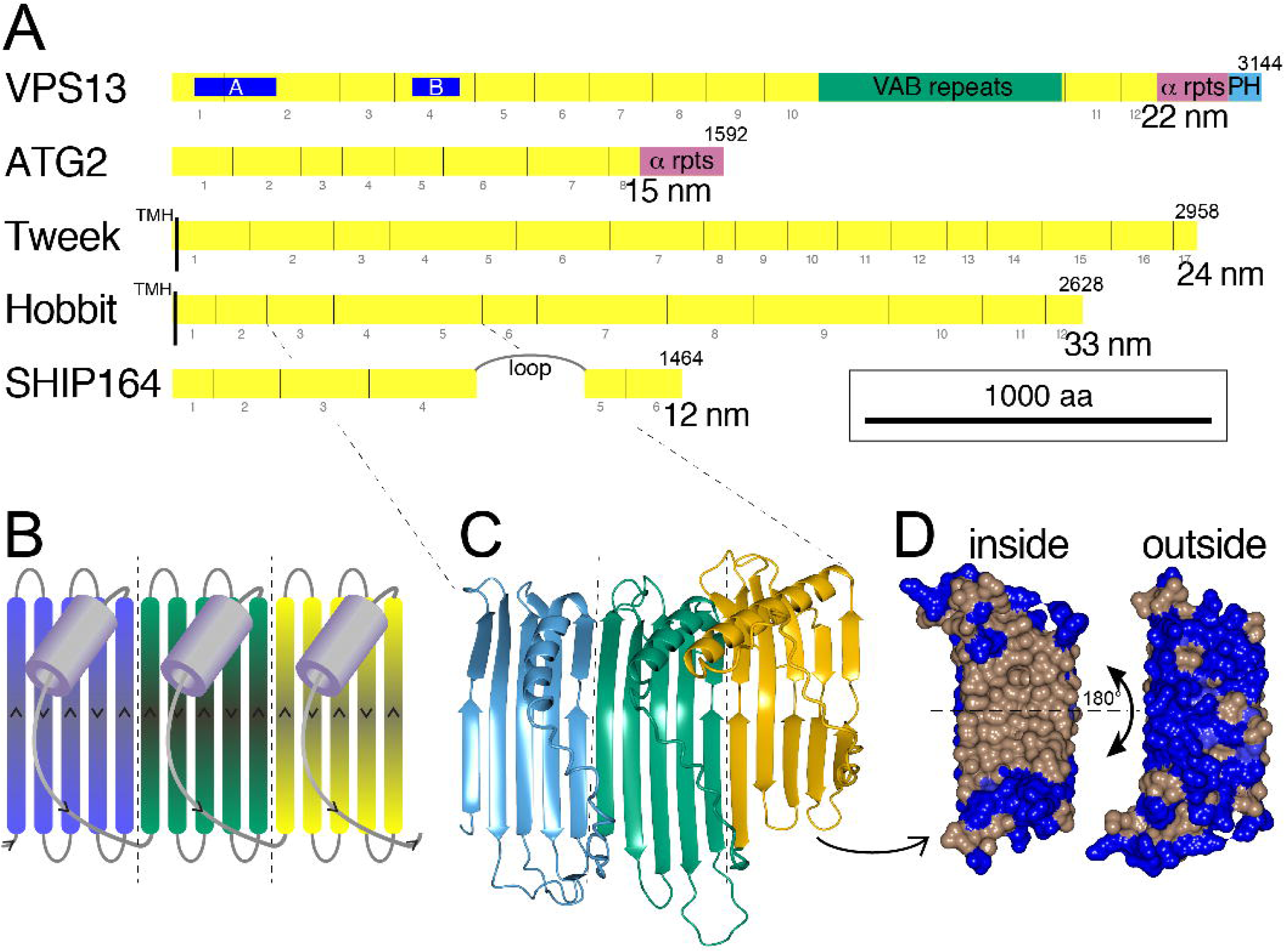
Five families of bridge-like lipid transfer proteins contain from 6 to 17 RBG domains. A. Domain structure in five families of eukaryotic bridge-like lipid transfer proteins. Repeating beta-groove (RBG) domains (yellow, also called “extended chorein-N domains” (Melia and Reinisch, 2022)) are numbered below. Also showing previously known domains with non-RBG structure: VAB repeats (VPS13 only, dark green), α repeats - tandem repeats (∼75 aa each) of paired helices in VPS13 and ATG2, also called ATG_C domains (magenta), pleckstrin homology (PH) domain in VPS13 (sky-blue), and transmembrane helices (TMH, black) (Tweek and Hobbit). An intrinsically disordered loop >300 residues long in SHIP164 is also indicated (Hanna *et al*., 2022), but loops shorter than 100 aa are not indicated. Blue regions A and B near the N-terminus of VPS13 (64-300 and 690-827) indicate bands where mutating hydrophobic sidechains lining the groove to charged residues (11 and 16 respectively) inhibited VPS13 function without altering the fold (Li *et al*., 2020). Predicted lengths of the RBG structures are as published elsewhere (Guillén-Samander *et al*., 2022; Neuman *et al*., 2022b). Homologs chosen are from yeast where possible (Vps13, Atg2, Csf1 and Hob1/Fmp27), as they are the most compact and have the smallest additional intrinsically disordered loops, or from human (SHIP164/UHRF1BP1L). B. Cartoon of three repeating β-groove (RBG) domains (colored blue/green/yellow in series). Each RBG domain consists of five β-strands in a meander pattern followed by the final sixth element consisting of an intrinsically disordered loop that most often starts with a short helix. Adapted from ref. (Neuman *et al*., 2022b). C. Structure of RBG domains 3, 4 and 5 from human Hobbit (KIAA0100, residues 272-729) predicted by AlphaFold, colored as in B. D. Surface of RBG5 (residues 569-695) showing the inside (concave) surface has hydrophobic side-chains (brown), while the outside (convex) surface is populated with hydrophilic side-chains (blue).

Analysis of the predicted structures of these proteins identified a repeating unit consisting of a 5-stranded β-sheet meander plus a sixth element that usually starts with a helix and then continues with a loop that crosses back over the meander (Figure 1B) (Neuman *et al*., 2022b). The rods are superhelical, which makes it hard to see the details of their construction (Guillén-Samander *et al*., 2022; Toulmay *et al*., 2022). There are some portions of predicted structure where the superhelical twist is low, and here the core building block is easily seen (Figure 1C). The individual unit of ≥150 residues has been named the Repeating Beta-Groove (RBG) domain (Neuman *et al*., 2022b). The concave (inside) surface of the β-sheet is hydrophobic and the convex (outside) surface is hydrophilic (Figure 1D). Because all 6 elements of βββββ-loop domain cross over the groove, the domain starts and ends on the same side of the groove, leading one domain to follow directly from another, repeating the same topology. Examination of the hydrophobic grooves of all five eukaryotic bridge-like LTPs shows that they are entirely assembled from multiple RBGs domains, with the rod-like proteins in each family being created from multimers of characteristic numbers of domains that determine length (Figure 1A) (Neuman *et al*., 2022b). Other bridge-like LTPs include those in the intermembrane spaces of mitochondria and chloroplasts. They more closely resemble their bacterial forebears (Neuman *et al*., 2022b) and are not considered here.

Here, the domain architecture of RBG superfamily is studied starting with whole proteins, moving to domains, and ending up with individual residues. Firstly, phylogeny, initially characterized for VPS13 decades ago (Velayos-Baeza *et al*., 2004), is updated indicating that the last eukaryotic common ancestor (LECA) had six RBG proteins: one from each family, and two VPS13 ancestors related to VPS13A/C/D and VPS13B. Secondly, the majority of central RBG domains arose by internal duplication from two complete domains in an ancestral protein with four domains. Thirdly, previously unknown accessory domains near the C-terminus of VPS13B and plant VPS13 homologs are described, which likely provide interaction sites for partners, alongside those already identified for membrane targeting (Bean *et al*., 2018; Kumar *et al*., 2018; Park and Neiman, 2020; Guillen-Samander *et al*., 2021) and for specific functions such as lipid scramblases (Ghanbarpour *et al*., 2021; Orii *et al*., 2021; Adlakha *et al*., 2022). Fourth, the extreme ends of RBG multimers are shown to all have amphipathic helices. However the C-terminus of SHIP164 is an exception as it has a coiled-coil. A hypothesis is developed that incorporates this and other bioinformatic evidence in a speculative model of SHIP164 function. Finally, conserved residues are distribute along the entire length of the external, hydrophilic surfaces of RBG proteins, indicating a greater number of sites for partner interactions directly with the bridge than previously envisaged.

## Results and Discussion

### A. Following RBG domains across evolution

#### (i) VPS13 forms a single unbroken groove

Prior to the AlphaFold predictions, it was known from low resolution cryo-EM studies that the LTP groove extended all the way along ATG2 (Valverde *et al*., 2019), and along a large portion of VPS13 (Li *et al*., 2020), but it was not clear whether the groove extended the full length of VPS13. AlphaFold predictions for ATG2, Hobbit, and SHIP164 show unbroken grooves running their full length (Jumper *et al*., 2021), but predictions for Tweek and for full-length VPS13 are missing, perhaps because the sequences are too long (Neuman *et al*., 2022b). Tweek has been constructed as one unbroken groove by overlapping partial models, made either by ColabFold, an online AlphaFold tool (Mirdita *et al*., 2021; Castro *et al*., 2022) or by trRosetta (Yang *et al*., 2020; Toulmay *et al*., 2022). In contrast, the first published full-length model of Vps13p made by overlapping fragments in showed RBG1-10 separate from RBG11/12 (Toulmay *et al*., 2022).

Linkage of the two segments of VPS13 was examined by a ColabFold prediction of the region encompassing both sides of the VAB repeats, with most of this central region being omitted. The prediction tool placed the two VPS13 segments in direct continuity (Figure 2B), with the final strand of RBG10 running parallel to the first strand of RBG11. The form of the RBG10/RBG11 interface precisely resembles that of any other RBG-RBG interface, for example RBG11/RBG12 (Figure 2C). Thus, ColabFold predicts that the rod-like molecule VPS13 forms a single continuous β-groove from end to end, resembling other members of the superfamily (Figure 2D). This matches cryo-EM observations of VPS13 as a rod (De *et al*., 2017) with a groove along at least part of its length (Li *et al*., 2020), and has been reported elsewhere (Adlakha *et al*., 2022; Guillén-Samander *et al*., 2022). This finding indicates that the VAB repeats, six all-beta domains of a unique type that build into a curved structure shaped like a hook (Bean *et al*., 2018; Adlakha *et al*., 2022), can be considered as a VPS13-specific insert in the loop between RBG domains (RBG10–11, see Figure 4A).

**Figure 2.**
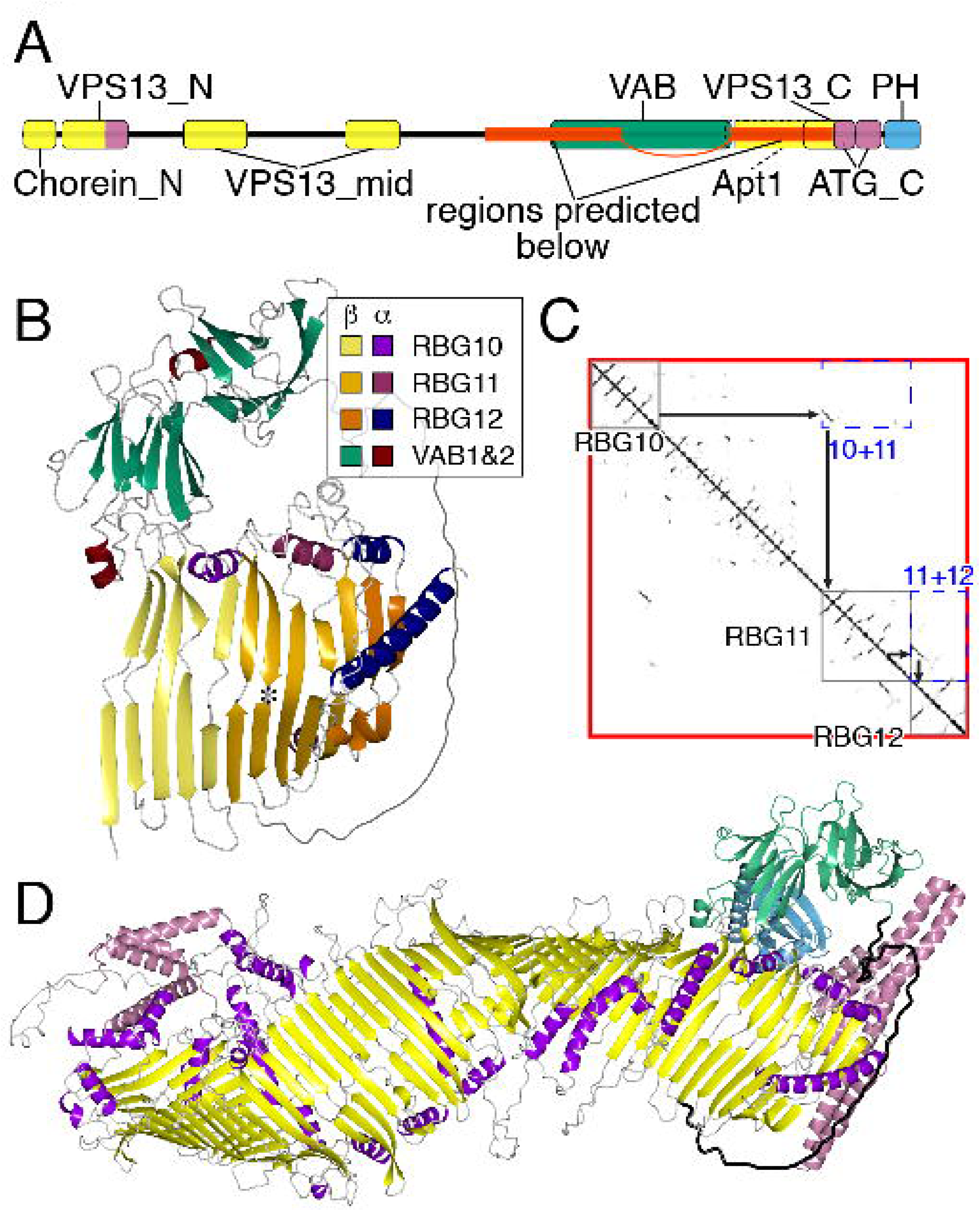
VPS13 makes one continuous hydrophobic groove. A. Domains of VPS13A identified in Pfam and by previous studies with HHpred, colored by underlying structure: (i) RBG domains (mostly β, yellow), which include: Chorein_N, VPS13_N, VPS13_mid, Apt1 (part) and VPS13_C; VAB repeats (all β, dark green) and the C-terminal pleckstrin homology (PH) domain, which is the only domain not unique to VPS13 (purple); (ii) all helical domains (magenta), which include: “handle” (C-terminus of VPS13_N, a four helical bundle containing the loops after both the RBG1 and RBG2 (Melia and Reinisch, 2022)) and ATG_C (overlaps C-terminus of VPS13_C). Two linked red sections indicate the region submitted to ColabFold (see B/C). “VPS13_N” refers to the domain called “VPS13” in Pfam (defined as “Vacuolar sorting-associated protein 13, N-terminal”), also called “VPS13_N2” at InterPro. Apt1 was originally defined in Hobbit and is homologous to the region indicated by dashed lines that includes RBG11/12 (Kaminska *et al*., 2016). B. Predicted structure for yeast Vps13 residues 1643-2111 + 2543-2840, which includes RBG10, VAB repeats 1&2, the unstructured linker that follows VAB repeat 6, and RBG11+12. The β/α portions of each domain/repeat are colored differently, see Key. Strand 5 of RBG10 is parallel to strand 1 of RBG11, in the same way that strand 5 of RBG11 is parallel to strand 1 of RBG12. The probability local distance difference test (pLDDT) of both RBG interfaces is ∼0.95. Asterisk indicates position in strand 2 of RBG11 where a WWE domain is found in VPS13A/C (see Section B). C. Contact map from AlphaFold, showing: internal interactions of each RBG domain near the main diagonal (grey boxes), off-diagonal interactions (RBG10+11, RBG11+12, blue dashed boxes), and predicted parallel contact by strand 5 of one RBG domain and strand 1 of the next domain (black arrows). D. Model of RBG multimer of VPS13, including the yeast Vps13 model the C-terminus as described in Methods. Regions are colored as in A, except: RBG domain helices are purple and disordered linkers are grey; the linker that follows the VAB repeats is highlighted in black. Note that no accessory domains extend beyond the end of the β-groove, with the PH domain lying behind the VAB repeats in this orientation.

Even though AlphaFold can reliably build 3D models from contact maps, the multimer predicted by AlphaFold varies from solved cryo-EM structure in more subtle aspects including both superhelicity and the extent to which the groove forms a sharp “V”, compared to a shallow groove (Li *et al*., 2020). Despite these limitations on AlphaFold (Adlakha *et al*., 2022; Neuman *et al*., 2022b), the VPS13 prediction has one possible biological implication, since it shows that the C-terminus of the lipid transfer groove in VPS13 is able to directly access a lipid bilayer, as the accessory domains do not project beyond the RBG multimer (Figure 2D). However, the links to the accessory domains are flexible, and other possibilities include that hydrophobic groove of VPS13 interacts with integral membrane proteins, for example scramblases (Ghanbarpour *et al*., 2021; Adlakha *et al*., 2022).

#### (ii) VPS13 has four major types of RBG domains

Sequence relationships between different RBG domains in VPS13 were examined to determine how the multimer of RBG domains was formed. The tool used for this was HHpred, which has high accuracy and a small number of known flaws (Fidler *et al*., 2016). A preliminary step was to identify all RBG domains in human VPS13 isoforms (Figure 3A). These do not align precisely with the Pfam domain structure, which has until now been the established to describe VPS13 structure (Figure 3A and 2A). VPS13A is the shortest both in terms of sequence length and in number of RBG domains: 12. By comparison, VPS13 and VPS13D have 15, and VPS13B has 13 (Figure 3A). All domains in VPS13 have five strands except RBG1 with 4 strands, the first being replaced by the N-terminal helix, and the final domain (RBG12 in VPS13A) with 2 strands (Neuman *et al*., 2022b). The 12 RBG domains in VPS13A align well with those in yeast Vps13 (yVps13) (Figure 3B) and in most plant homologs (Figure 3C), indicating that this is an ancient form. The increased number of domains in VPS13C and VPS13D fits with previous observations that their extended length originated from internal duplications of ∼500 residues (Kumar *et al*., 2018; Guillen-Samander *et al*., 2021). The basis for variation in RBG domain number may relate to width of contact size bridged by individual homologs in a way that has yet to be studied.

**Figure 3:**
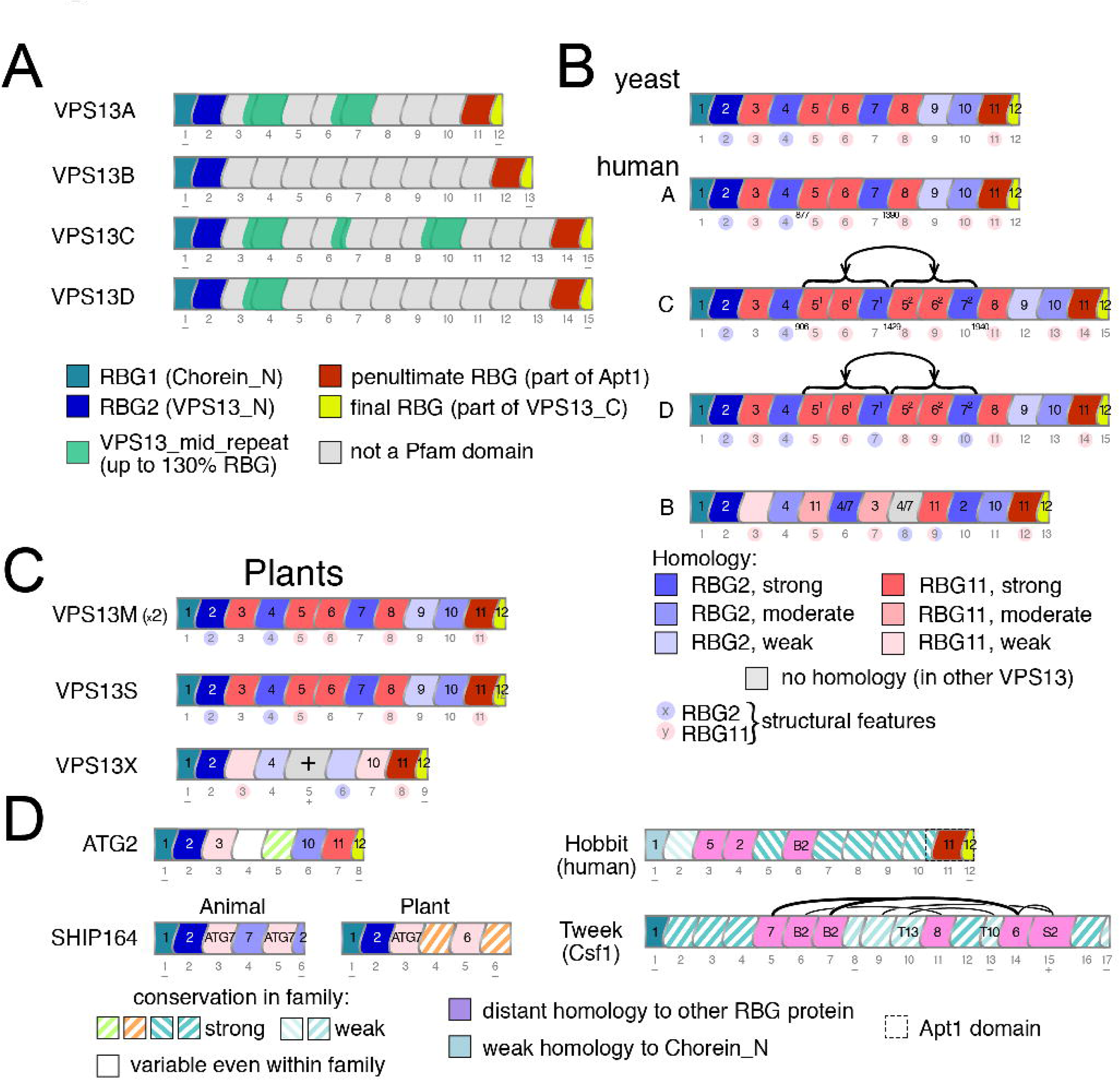
Homology between RBG domains indicates patterns of inheritance from shorter ancestral forms. A. Schematic representation of RBG domains in VPS13A–D, including their mapping to Pfam domains, colored as in the key. RBG domains are drawn in a stylized way to indicate that loops do not cross the β-groove perpendicularly; width of domains is determined by the number of β-strands: typically 5, but RBG1 and RBG12 narrower (4 and 2 strands respectively). Here, and in B–D following, the position of each RBG domain is indicated by numbers below, and “+”/“–” indicate variations from 5 strands per domain. B. Homology relationships to RBG2 (blue) or RBG11 (red) for central RBG domains in yeast Vps13 and human VPS13A–D. Depth of shading indicates stronger homology as in the key below. Numbers inside domains indicate the orthologous RBG domain in VPS13A, except VPS13B- RBG3, which is has no homology to VPS13A, and is weakly similar to RBG12 in VPS13B (indicated by light red). Duplications indicated by bracketed areas, with superscript^1/2^ for the numbers inside domains, and residue numbers for the duplication boundaries in VPS13A/C. Minor structural variations similar to RBG2 (loop) or RBG11 (bulge), or both (see Section Dii) are indicated by light blue or light red circles around domain positions (also in C). C. Homology relationships as in B for plant VPS13 proteins: VPS13M1/2 (highly similar paralogs, only one shown), VPS13S and VPS13X. RBG5 in the latter has no discernible homology outside plant orthologs. D. Homology relations of RBG domains in ATG2 (human ATG2B), SHIP164 (human and *Arabidopsis*), Hobbit (human) and Tweek (yeast Csf1), domain boundaries as in Supplementary File 1. Diagram summarizes findings from diverse species, except for SHIP164 where animal and plant proteins are shown separately. ATG2 RBG4 is the only domain showing no significant conservation across evolution (filled white). Homology with domains in VPS13A/VPS13B/ ATG2/SHIP164/Tweek and numbers inside prefixed by nil/B/ATG/S/T respectively. Hatching in domains indicates moderate or strong homology only within the same RBG family, for example for Hobbit: it indicates homology to RBG domains beyond the animal kingdom. The depth of color correlates with the degree of conservation. Homology between Tweek domains is shown by lines above, thickness correlating strength of homology.

The next step was to identify homologies between individual RBG domains. When searches are seeded with whole protein strongly homologous regions in some hits will cause false positive alignments of adjacent unrelated regions (Fidler *et al*., 2016). This is a particular problem in proteins with repeats (data not shown). Therefore, multiple sequence alignments (MSAs) were created for each RBG domain separately using 5 iterations of HHblits to search deeply for homologs across human, yeast, protist and plant proteomes. In these searches, all RBG domains produced strong hits to the orthologous region of VPS13 in model species across a wide range of eukaryotic evolution (probability of shared structure >98% in fly, worm, yeast, *Capsaspora, Trichomonas, Trypanosoma, Chlamydomonas* and *Arabidopsis*). One aspect of the method that was not based on biological observation is that the six elements of RBG domains were defined as βββββ-loop. This was chosen in preference to any other permutation (for example βββ-loop-ββ) to maximize the power of sequence analysis, because either including the whole loop that follows the helix, the site of greatest variability, or making a deletion reduces the sensitivity of MSAs (not shown). Indication that RBG domains may not form the βββββ-loop biological unit are highlighted in the descriptions of search results below.

Searches based on each RBG domain showed that all domains have strong hits to orthologous regions, which can therefore be considered as RBG domain orthologs. A new observation that the central 10 domains of VPS13A (RBG2 to -11) all show homology to other domains. This was seen in a second tranche of hits (typically with probability of shared structure up to 95%, and down to 10%) aligning each domain to other regions of VPS13 (and in some cases to ATG2 and SHIP164). This indicated more distant relationships between RBG domains than have been described approximately before (Kumar *et al*., 2018; Castro *et al*., 2022). To establish relationships while avoiding all-vs-all comparisons, the situation was simplified by noting that domains tend to fall into one of two types, either similar to RBG2–the penultimate domain at the N-terminus RBG2, or similar to RBG11–the penultimate domain at the C-terminus (Figure 3B). Some of these relationships are also supported by clustering of RBG domains solely using BLAST, for example clustering RBG4 both with RBG2 and RBG7 (Supplementary Figure 1). Eight out of ten domains showed strong homology (predicted shared structure >90%) to one but not both of the N-terminal or C-terminal domain (Supplementary Table 1A). The two domains not in this group were RBG10 with moderate homology (predicted shared structure 77%), and RBG9 with weak and mixed homology (predicted shared structure 40% for N-terminus and 18% for C-terminus) (Supplementary Table 1A).

The same findings were made for VPS13C and VPS13D, except three of the central domains of both (RBG5–7) had very close homologs 3 domains distant in the multimer (Figure 3B, see numbers 5^1^–7^1^ and 5^2^–7^2^ inside domains). The inferred duplication events are located at position nearer the N-terminus than previously suggested (Kumar *et al*., 2018). The origin of the extra sequence in VPS13C/D was confirmed by examining sequence relationships between >200 RBG domains from VPS13 homologs in 7 model organisms. A cluster map confirmed close relationships between the VPS13C pairs RBG5^1+2^, RBG6^1+2^, and RBG7^1+2^ (Supplementary Figure 1). RBG5^1+2^, RBG6^1+2^, and RBG7^1+2^ from VPS13D were also homologous to each other, but much more weakly. This suggests that RBG5/6/7 duplicated in two independent events, more recently in VPS13C than in VPS13D.

Overall, this shows that VPS13A/C/D in their full extent consist of domains related to RBG1-2-11-12, which are therefore are the four major types of RBG domain in VPS13. In turn this suggests that a very primitive ancestor of VPS13 may have consisted of just these four domains, with growth by repeated central duplication. Appearance of new domains centrally like this is the typical pattern of duplication of multimeric domains in one protein (Bjorklund *et al*., 2006). While divergence for RBG3 to -8 is limited (homology is strong), divergence has been greater for RBG10 and RBG9. The mixed homology in the latter is not unique (Supplementary Table 1A). Here opposite ends of the RBG domain are homologous to RBG2 and RBG11 (data not shown), which is evidence for inheritance of RBG domains in units other than βββββ-loop. Features that differentiate between RBG2 and RBG11 in terms of both structure and sequence are described in Section D.

#### (iii) VPS13B diverged from VPS13A/C/D at the 6+ RBG domain stage

VPS13B is an outlier compared to the others in its VAB repeats (Dziurdzik *et al*., 2020), and the same applies for its RBG domains. Six RBG domains (RBG1/2/4/11/12/13) are close in sequence terms to VPS13A (RBG1/2/4/10/11/12, with a 7^th^ domain is well conserved but not in sequence (RBG7 in VPS13B is similar to RBG3 in VPS13A). The remaining RBG domains in VPS13B follow a quite different pattern (Figure 3C). Five of these, RBG5/6/8/9/10, do not align well with any of the domains of VPS13A/C/D in the same position, instead resembling a mixture of domains or a penultimate domain. In addition, one domain (RBG3) has just weak and partial homology to the C-terminal penultimate domain and resembles no other domain specifically (Figure 3C and Supplementary Table 1A). These findings indicate that VPB13B started to diverge from VPS13A/C/D at or after a 7 domain stage consisting of RBG1-2-3-4-10-11-12 (numbering according to VPS13A). The timing of this divergence is addressed in the next section.

#### (iv) Phylogeny of VPS13A/C/D and VPS13B indicates LECA expressed these two isoforms

Looking across evolution to construct a phylogeny for VPS13, in invertebrates the fruit fly *D. melanogaster* has three VPS13s: clear homologs of VPS13B and VP13D, plus one protein called *Vps13* that is related to both VPS13A and VPS13C. This is consistent with the VPS13A/C pair arising from a relatively recent duplication (Ugur *et al*., 2020). The domain structures of the three fly VPS13s are identical to human VPS13A/B/D, suggesting that VPS13C is the divergent vertebrate homolog. Among other invertebrate model organisms, the nematode worm *C. elegans* has two VPS13s that resemble VPS13A/C (gene: T08G11) and VPS13D (C25H3.11) (Brickner and Fuller, 1997; Velayos-Baeza *et al*., 2004). VPS13D in *C. elegans* (and in other worms, not shown) has eleven RBG domains: in detail it lacks RBG3 and one set of RBG5-6-7. It also does not contain the UBA domain found in human and fly VPS13D (Anding *et al*., 2018; Shen *et al*., 2021). Other invertebrates were examined to provide context for the situation in worms. The simple eukaryote *Trichoplax adherens* also has the VPS13A/C and VPS13D pair, and *T. adherens* VPS13D has the same 15 RBG domains as human, which indicates that worms are outliers in their loss of domains. For VPS13B, even though it is missing from *C. elegans* and *Trichoplax*, a wider search showed that some invertebrates have VPS13B.

These findings indicate that RBG domains are both gained and lost across evolution, but does not address how ancient the form VPS13B is. To determine if ancestral versions of VPS13B and VPS13D predate the evolution of animals, more divergent genomes were examined. *Capsaspora owczarzaki*, a free-living single cell organism related to animal precursors (Suga *et al*., 2013) has four VPS13s, three of which resemble VPS13A/C, VPS13B and VPS13D in flies, indicating that VPS13B and D both originated before animals evolved. Looking deeper into evolution, the slime mold *Dictyostelium discoideum*, which diverged from the common animal/fungal ancestor, has multiple VPS13s (Leiba *et al*., 2017), but none are related to either VPS13B or VPS13D, and this is the case in all amoebae (not shown). To determine if the absence of VPS13B and VPS13D in amoebae is because they evolved in opisthokonts only, VPS13 sequences were compared across the whole of eukaryotic evolution. In a cluster map of >1000 proteins, VPS13D was closely linked to VPS13A/C, while VPS13B clustered separately (Supplementary Figure 2). A key finding is that 8% of the VPS13B cluster were from SAR/Harosa protists and algae (Supplementary Figure 2), with species such as *Aphanomyces* containing full-length homologs of VPS13B (data not shown). By comparison, the VPS13D cluster contained one amoebal protein and one algal protein, and the basis for such clustering was unclear as each showed stronger BLAST hits to VPS13A/C than to VPS13D. Thus, in agreement with the RBG domain results (Figure 3B), it appears that VPS13D split from VPS13A/C in pre-metazoal evolution, and that VPS13B is an ancient paralog of VPS13A/C/D so widespread that it is likely to have also been in LECA. This conclusion differs from prior work that that related VPS13B to one specific plant homolog (Velayos-Baeza *et al*., 2004) or to VPS13D (Leiba *et al*., 2017), possibly because those assignments relied on proteins from a small number of model species rather than as large a range of protein sequences as possible.

While the analysis here is based on assigning orthology, VPS13 in protists separated from other extant organisms by long evolutionary branches cannot be assigned to any one group. They still merit study as they are examples of the general principle of plasticity in RBG proteins. Individual VPS13 proteins in *Chlamydomonas Plasmodium*, and *Toxoplasma* have 7192, 9307 and 13455 amino acids respectively, the latter containing 188 predicted β- strands (not shown), suggesting expansion to 38 RBG domains with a hydrophobic groove >75 nm. It may be that intracellular parasites have unique inter-membrane contacts that are wider than typical eukaryotic cells.

#### (v) VPS13X is a previously undescribed divergent plant homolog

Until now, plants have been thought to contain three VPS13 homologs (Velayos-Baeza *et al*., 2004). To describe these briefly, the only plant VPS13 that has been studied experimentally is *SHBY*(At5g24740), named because mutations cause *Arabidopsis* appear **<u>sh</u>**rub**<u>by</u>** (Koizumi and Gallagher, 2013). The name VPS13S (from ***<u>S</u>HBY***) is proposed here to standardize the format across VSP13 proteins in major clades of organism where possible. The two other identified *Arabidopsis* VPS13 proteins (At4g17140 and At1g48090 (Velayos-Baeza *et al*., 2004)) have not been named or studied directly. They are close paralogs and the contain multiple accessory domains (described in Section Biii), so the names VPS13M1/2 (for **<u>m</u>**ultiple) are used here. VPS13S and VPS13M1/2 have RBG domains homologous to VPS13A and yVps13 (Figure 3C), consistent with this being the arrangement of RBG domains in LECA’s VPS13A/C/D ancestor.

HHpred identified a fourth VPS13 in *Arabidopsis* (At3g50380), which is annotated in databases as “Vacuolar protein sorting-associated protein”, named here VPS13X and has close homologs in most land plants (data not shown). VAB repeats two domains from the C-terminus identify it definitively as a VPS13 rather than ATG2, but it is variant being shorter than other VPS13s with nine RBG domains only six of which are homologous to domains in other VPS13s: RBG1-2-4-7-8-9 follow the same pattern as RBG1-2-4-10-11-12 (VPS13A numbering, Figure 3C). Two domains (RBG3 and -6) are unrelated to specific VPS13 domains, though they have distant relationships to the penultimate domains (Supplementary Table 1A). Finally, one domain (RBG5) is homologous only to the orthologous domain in other VPS13X proteins, and has seven β-strands, confirmed by ColabFold (Supplementary Figure 3). This high level of divergence makes it impossible to tell if VPS13X originated from VPS13A/C/D or from VPS13B or from their common ancestor.

#### (vi) ATG2 shares six RBG domains with VPS13

Relationships between the eight RBG domains of ATG2 were traced as had been done for VPS13 (Supplementary Table 1B, Figure 3D). The four domains nearest the ends (RBG-1/2/7/8) are strongly homologous to the four major RBG types in VPS13A, which adopt equivalent positions (RBG-1/2/11/12). The next pair inwards show homology to domains RBG3/10 in VPS13A, though weak for RBG3. Finally, the most central two domains (RBG4 and -5) cannot be traced to any domain outside ATG2 itself. While one (RBG5) is well conserved among ATG2 proteins, the other (RBG4) is highly variable across evolution, for example *S. cerevisiae* and *S. pombe* domains being unrelated. These results suggest that ATG2 and VPS13 have a common ancestor with 6 RBG domains, thus possibly preceding the split in VPS13 at a 7 domain stage.

#### (vii) SHIP164 lacks a specialized C-terminus

SHIP164 has homologs from animals to plants with six RBG domains, with homology confined to the their N-termini (RBG1–2), resembling the N-termini of both VPS13 and ATG2 (Figure 3D). For RBG3, a common ancestor related to RBG7 of ATG2 can be detected for both animals and plants, but they have diverged so far as to share no homology. In the C-terminus, there are some domains related to others outside the family (Supplementary Table 1C), but RBG4 and RBG6 in plants are unique forms, and the latter is considerably different from the same domain in animals, having 5 strands and 2½ strands respectively. The homology of the final domain of SHIP164 in animals to RBG2 of VPS13 is significant because it groups the domain away from those specialized to be at the end of the multimer, suggesting that the C-terminus is similar to the middle of an RBG multimer (see Section C).

#### (viii) Hob and Tweek proteins show more distant homologies to VPS13

Before mapping RBG domains in these proteins, a preliminary step was to check previous reports that homologs of Tweek exist only in fungi (Csf1) and animals, such as fly (Tweek) and human (FSA/KIAA1109) (Neuman *et al*., 2022b; Toulmay *et al*., 2022). Using HHpred, full-length homologs were identified in *Trypanosoma* (4203 aa, XP_011775923) and *Trichomonas* (2695 aa, XP_001306426), lengths within the range between yeast Csf1 (2958 aa) and fly Tweek (5075 aa) or human KIAA1109 (5093 aa). All length variation in this family results from intrinsically disordered loops throughout the proteins. Although horizontal gene transfer into multiple protists cannot be excluded, this suggests that Tweek/Csf1 was present in LECA, which is a more ancient origin than was previously considered.

Hobbit and Tweek both start with transmembrane helices (TMHs) that anchor them in the ER (Figure 1A) (Castro *et al*., 2022; Hanna *et al*., 2022; John Peter *et al*., 2022; Toulmay *et al*., 2022). A feature at the N-terminus of Hobbit and Tweek is that their RBG2s are dissimilar to the RBG2 shared by VPS13, ATG2 and SHIP164. However, both Hobbit and Tweek have domains that are distantly related to VPS13/ATG2/SHIP164 RBG2 (RBG4/6 and RBG6/7/15 respectively), indicating that this domain was present in their ancestral forms (Figure 3D). The lack of RBG2 in its standard position might reflect reduced evolutionary pressure for protein interactions at the N-terminus compared to RBG domains because of the TMHs, but other domains are distantly related to RBG2. The other three major RBG domain types (RBG1/11/12, VPS13 numbering) also have homologs in Hobbit. The strong C-terminal homology has been noted previously (Rzepnikowska *et al*., 2017), while the weak N-terminal homology is a new finding. This leaves 6 other domains conserved within Hobbit family (Figure 3D). In Tweek, only two of the major RBG domains can be found: RBG1 is strongly homologous to RBG1 in VPS13/ATG2/SHIP164, and three domains are weakly similar to RBG2. The major RBG domains types at the C-terminus (RBG11/12 in VPS13) are not found, similar to SHIP164. Three other Tweek domains (RBG5/11/14) show weak homologies to three central VPS13 domains (6/7/8), a different set of domains than occurs in Hobbit. This suggests that Hobbit and Tweek both have common ancestors with the other RBG proteins, but the precise form cannot be determined. Relationships between two groups of Tweek domains (RBG5–10 and RBG11–15) suggest an internal duplication. All domains in Tweek are conserved across the family, and three have variant numbers of β- strands (3, 3 and 7 β-strands in RBG8/10/15) conserved from human to *Trichomonas*. This indicates that acquisition of domains and their rearrangement all occurred before LECA.

To summarize the whole section on conservation: the majority of RBG domains have conserved sequence across all eukaryotes. A minority of RBG domains (VPS13X RBG5, ATG2 RBG4, Hobbit RBG4) have changed so much that the individual RBG domain is not detectable across all orthologs. This minority shows that primary structure can change to lose detectable homology without affecting secondary structure or lipid transfer function. In turn this implies that the conservation present in all the other parts of RBG proteins serves specific functions other than lipid transfer itself. Section D addresses these conserved sequences in detail in order to better understand their functional role.

### B. The range of accessory domains is wider than previously known, providing more ways to interact with partners

The widely conserved accessory domains are a set of three near the C-terminus of VPS13 (VAB, ATG_C and PH), ATG_C also being found in ATG2. After delineating the RBG domains, it became possible to identify all other accessory, folded, non-RBG domains using HHpred with confirmatory modelling by ColabFold (Figure 4A). Being able to benchmark HHpred against AlphaFold predictions simplifies recognition of variant RBG domains, even if they contain multiple and extended inserts of small helices and disordered loops, or part of their secondary sheet structure is mistakenly identified as helix by PSIPRED, the tool used by HHpred (Buchan *et al*., 2013). Below, additional alpha helical domains are discussed first, and then a range of mainly-beta domain inserts that are far more numerous among VPS13 homologs than previously thought.

**Figure 4:**
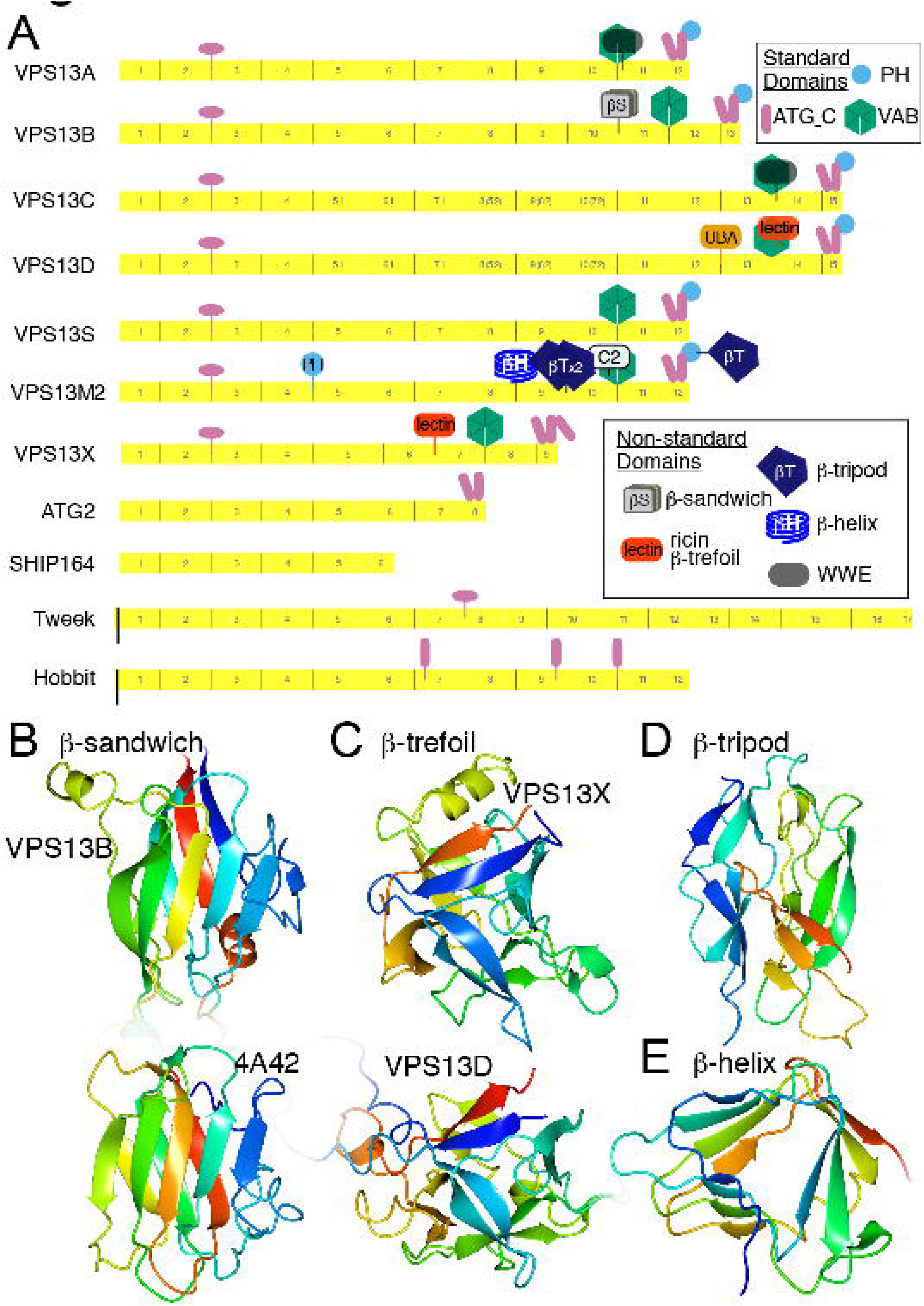
VPS13 homologs, including VPS13B, have accessory domains in greater number than previously known A. Full domain structure of RBG proteins, including all human and *Arabidopsis* VPS13 homologs (except VPS13M1, which is same as VPS13M2 but missing the C-terminal β-tripod). Domains include: transmembrane helices (TMH – black bars), α-bundle (pink ovals, including “handle” in VPS13), VAB repeats (green hexagon), anti-parallel paired helices (called ATG_C, pink cylinders), PH (purple circle), previously well-known accessory domains: UBA, ricin-like lectin (β-trefoil) in VPS13D, PH (N-terminal) and C2 as indicated. Additional domains: <u>all-α:</u> bundle; <u>mixed α/β:</u> WWE; <u>mostly−β:</u> ricin-like lectin in VPS13X, βH=β-helix, βS=β-sandwich, βT=β-tripod (see key). Despite the β-tripods being called VPS62 in Pfam, the predicted structure aligns with the Gram-negative insecticidal protein 6fbm (z=15.8, RMSD 2.9Å for 166 residues); and the AlphaFold model of Vps62 lacks the fold (data not shown); additionally the β-tripod–VPS62 link has been falsified (Zaitseva *et al*., 2019). B. Structure of β-sandwich insert in VPS13B RBG10/11 loop predicted by ColabFold, which aligns with discoidin domains and similar carbohydrate binding domains (z=8.1, rmsd 3.1 Å, across 112 residues), here showing below for comparison a carbohydrate-binding module-32 (CBM32) domain from a *C. perfringens* alpha-N-acetylglucosaminidase family protein (PDB: 4A42). C. Structure predicted by ColabFold of the insert between RBG6 and RBG7 of VPS13X (1426-1585). DALI aligns this to haemagglutinin (PDB: 4OUJ, z=14.8 rmsd=2.1 Å, across 117 residues), confirming HHpred’s probability shared structure (pSS) =88% with ricin-type β-trefoil lectins. The structurally similar domain in VPS13D (3592-3780) is shown for comparison below. D. VPS13M1 residues 2262-2421 predicted by ColabFold as a β-tripod, which appears as a tandem pair, plus an extra copy at the C-terminus of VPS13M2. E. ColabFold prediction of VPS13M1 residues 1726-1840, forming a 4-turn β-helix. In B–E, structures are colored in a spectrum from N-terminus (blue) to C-terminus (red). pLDDT of the models of new domains (excluding inserted loops) in VPS13B, VPS13X, and the β-tripod and β-helix in VPS13M was 0.88 over 142 residues, 0.89 over 136 residues, 0.88 over 158 residues and 0.95 over 110 residues (B–E respectively).

#### (i) Helical bundles similar to those in VPS13 and ATG2 are found in Hobbit in Tweek

The best known accessory domain in any VPS13 is the ∼45 aa helical UBA domain in VPS13D (Velayos-Baeza *et al*., 2004). This is inserted in the RBG12/13 loop (Figure 4A), and may be related to VPS13D’s role in mitophagy, given its typical role in binding polyubiquitin (Anding *et al*., 2018; Shen *et al*., 2021). UBA domains also occur at the extreme C-terminus in a minority of some algal VPS13 homologs, indicating a strong selection pressure for UBA domains in this family. RBG proteins have only a few other helical regions, one of which has solved structure: the VPS13 “handle”, a helical bundle with unknown function near the N-terminus where extra helices occur in 2 loops (RBG1/2 and RBG2/3) (Li *et al*., 2020). Secondary and tertiary structure of other helical regions among the RBG proteins were predicted by HHpred and AlphaFold respectively. The best known helical region is the ATG_C region (∼150 aa) in ATG2 that targets lipid droplets and autophagosomes and is amphipathic (Figure 4A) (Velikkakath *et al*., 2012; Kotani *et al*., 2018). VPS13 has a homologous region that targets lipid droplets (Kumar *et al*., 2018), and a homologous region is present in a subgroup of fungal sterol glucosyltransferases (Grille *et al*., 2010). Alphafold suggests that the whole ATG_C region contains four main helices forming two repeats of an anti-parallel helical pair (∼75 aa). Although AlphaFold orients the two bundles at a specific angle to each other, with a low probability of local distance difference test (pLDDT) for this region, suggesting the bundles are highly mobile, with no support for any particular relative orientation. Alphafold predicts three similar anti-parallel helical pairs in Hobbit (two conserved in plants), but none in SHIP164 or Tweek, the latter having a conserved helical bundle (120 residues) near its middle (Figure 4A). Each helical region contains at least one amphipathic region ≥18 residues, as used for membrane targeting by ATG_C (Kumar *et al*., 2018), however the regions in Hobbit and Tweek are not located near the end of the Hobbit and Tweek bridges, so it is not clear how they might interact with membranes when the protein is bridging between membranes, but they might act at other times.

#### (ii) VPS13B contains a carbohydrate binding accessory domain, matching the function of the lectin domain in VPS13D

Other accessory domains that have been described before, though not yet studied, include a second domain in VPS13D and one domain each in VPS13A and VPS13C. The other domain in VPS13D is a Ricin-type β-trefoil lectin inserted in a loop of the sixth VAB repeat (Figure 4A) (Velayos-Baeza *et al*., 2004), which may indicate a role in binding an O-linked GlcNAc group that can be reversibly added to serines and threonines of cytoplasmic proteins particularly in nutrient and stress sensing pathways, including regulators of autophagy (Hart *et al*., 2007,Hart, 2019 #3137). The domains in VPS13A/C are 75 α/β WWE domains inserted in RBG11, although databases annotate only a minority of cases (Figures 7A) (Cai *et al*., 2022; Hanna *et al*., 2022). WWE domains are thought to bind proteins involved in either ubiquitination or poly-ADP-ribosylation (Li and Chen, 2014). Together with the UBA domain of VPS13D, this may reflect consistent pressure for VPS13 proteins to interact with the ubiquitination machinery.

While VPS13B was not previously thought to have an additional domain, the survey here revealed a 180 residue mostly-beta domain inserted in the RBG10/11 loop (Figure 4A). This might have been missed previously because it has no sequence homologs, and its mostly-β structure is hard to distinguish from the typical pattern of RBG domain elements. HHpred identified the domain in all VPS13B homologs, including in protists (*e.g.* the oomycete *Phytophthora*). ColabFold modelled this as a β-sandwich (Figure 4B). The same β-sandwich is found in the discoidin domain, which binds carbohydrate, so VPS13B and VPS13D potentially have accessory domains with similar functions, although the structure and position differs from the lectin in VPS13D.

#### (iii) Multiple accessory domains link plant VPS13 proteins to ubiquitination

Plant homologs vary considerably in their accessory domains. While VPS13S is the same as yVps13, VPS13X is the only protein studied here that lacks any of the characteristic accessory domains: its C-terminal PH domain is replaced by three helices with amphipathic properties, possibly extending the ATG_C region. VPS13X also contains an additional 145 residue mostly- β domain in the loop between RBG6 and RBG7. ColabFold predicts this to be a Ricin-type lectin, with the same typical β-trefoil structure found in VPS13D, though lacking any sequence homology (Figure 4C) (Parker *et al*., 2021). The presence of functionally identical domains in VPS13X (plants) and VPS13D (opisthokonts) might indicate an ancient relationship between these proteins, or repeated acquisition of the same accessory domain.

In contrast to other VPS13 homologs with one extra domain, VPS13M1/2 have many (6 and 5 respectively, Figure 4A). In Pfam VPS13M1/2 are identified as respectively containing a PH domain in the RBG4/5 loop or a C2 domain inserted the first loop of the first VAB repeat. HHpred extends this to discover PH and C2 domains in both VSP13M1 and -M2. Pfam also documents a tandem pair of “Vps62 domains” in both proteins, while HHpred finds a third such domain in a loop of the C-terminal PH domain of VPS13M1 only. AlphaFold predicts these domains as β-tripods (Figure 4D), a structure first described in bacterial lysins, where a possible function in protein binding has been identified, and the confusion with Vps62 is explained as a spurious alignment error (Zaitseva *et al*., 2019). Finally, HHpred identified a four-turn right-handed β-helix located in the RBG8/9 loop of VPS13M1/2 (Figure 4E). This domain is also found in some protist VPS13s (not shown), and previously was found in various proteins including F-box proteins such as FBXO11 (Yoder *et al*., 1993; Ciccarelli *et al*., 2002). Thus, the analysis of domains in plants suggests yet another link for VPS13 to ubiquitination pathways.

Overall, all four human VPS13s have non-standard accessory non-RBG domains near the C-terminus and an even more complex situation has evolved in plants (Figure 4A). Knowing the location of these domains adds to the catalog of sites likely to bind interaction partners, as already known for VAB and PH domains (Bean *et al*., 2018; Dziurdzik *et al*., 2020; Park and Neiman, 2020; Guillén-Samander *et al*., 2022). In yeast just one homolog carries out the multiple functions of VPS13, and correspondingly yVps13 has multiple intracellular locations (De *et al*., 2017; Dziurdzik and Conibear, 2021). It is possible that the many different accessory domains facilitate the division of VPS13 function in complex organisms into subsets that are regulated independently.

### C. Extreme ends of RBG multimers have amphipathic helices – with the exception of the C-terminus of SHIP164

If RBG proteins bridge across membrane contact sites, each extreme end might interact with a bilayer to allow lipid entry/exit. This section looks at the properties of structural elements at the ends of RBG multimers, focussing on a common finding that they are capped by amphipathic helices, and an exception at the C-terminus of SHIP164, which has a different kind of helix, a coiled-coil.

#### (i) RBG multimers without TMHs start with an amphipathic helix, except ATG2

At the N-terminus there is uniformity across the entire RBG superfamily, with all RBG1 domains being homologs of the Chorein_N domain, even though for Hobbit this homology is weak (Figure 3). In one study of targeting of ATG2, just 46 residues at the N-terminus were needed for ER localization (Kotani *et al*., 2018), which includes just the start of RBG1/Chorein_N. RBG1 uniquely has the first strand replaced by a helix that crosses the multimer perpendicularly, in contrast to the helices that occur in the middle of the multimer that align along it (see Figure 7). To look for adaptation for interaction with target bilayers, these helices were examined for potential amphipathicity (Gimenez-Andres *et al*., 2018). VPS13 starts with a helix with amphipathic properties that are well conserved, being on average uncharged and having a broad hydrophobic face (9 residues, Figure 5A and Supplementary Table 2A).

**Figure 5:**
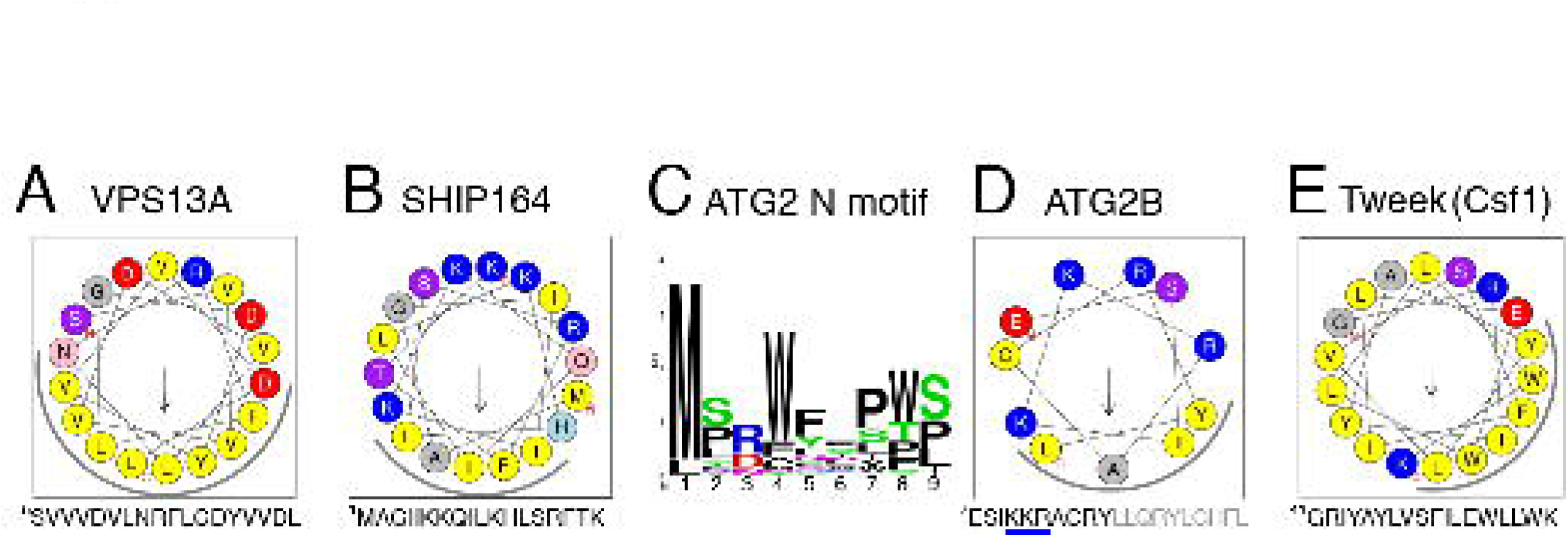
Features at the extreme N-termini of RBG proteins. A. Helical wheel projections (from Heliquest) of residues 5-22 of VSP13A, created by Heliquest, with hydrophobic face indicated by an arc around the lower surface. B. Amphipathic helix in SHIP164 (1-18), as in A. C. Peptide consensus of the segment prior to initial helices of ATG2 N-termini from 46 sequences across evolution; median length 8 (6- 15) residues with conserved large hydrophobic side-chains. D. Helical wheel projection of the first 11 residues only of the initial helix of human ATGB (7-17), with remaining helix indicated by grey letters below. Three positive residues ^10^KKR^12^ in ATG2B that are homologous to ^10^QKR^12^ in yeast Atg2 that mediate ER targeting by an unknown mechanism are underlined in blue (Kotani *et al*., 2018). E. Helical wheel projections of the transmembrane helix (TMH) and amphipathic helix (AH) from the yeast homologs of Tweek (Csf1), showing the hydrophobic face as in A.

One possibility is that these amphipathic helices interact with protein partners, such as the his is consistent with insertion in a membrane with many packing defects and low levels of anionic headgroups, such as the ER where the N-terminus targets in almost all examples studied of VPS13 (Kumar *et al*., 2018, Gimenez-Andres, 2018 #2906,Gonzalez Montoro, 2018 #3152). Exceptions include VPS13B with a +3 charge, which might explain its ability to “moonlight” on endosomes (Koike and Jahn, 2019). Even more extreme, the amphipathic helices in SHIP164 and VPS13X have narrow hydrophobic faces and are highly charged (Figure 5B, Supplementary Table 2A), which is consistent with these N-termini inserting into tightly packed membranes with high levels of anionic headgroups, particularly the plasma membrane (Bigay and Antonny, 2012; Hanna *et al*., 2022).

ATG2 differs from VPS13 and SHIP164 by commencing with an unstructured hydrophobic loop that can enhance membrane insertion (Figure 5C) (Fuglebakk and Reuter, 2018) and distantly resembles the “WPP motif” that targets plant proteins to the nuclear envelope (Patel *et al*., 2004). The following helix has a short amphipathic section (11-13 residues) (Figure 5D). This helix is important for ER targeting, as it includes three charged residues required for Atg2 function (Kotani *et al*., 2018). Although it is unknown how these two elements for membrane interaction might function together, they might account for the ability of ATG2 to transfer lipids from more than one source (Noda, 2021), particularly highly curved tubules (Maeda *et al*., 2019).

The N-termini of both Hobbit and Tweek start with TMHs that integrate into the ER (Figure 1A) (John Peter *et al*., 2022; Neuman *et al*., 2022a; Toulmay *et al*., 2022). While Hobbit has only a TMH, in Tweek and homologs the TMH is followed by an amphipathic helix (Figure 5E and Supplementary Table 2A). Since this is unlikely to dominate over the TMH in ER targeting, its role might be to select regions of curvature and/or to locally disorganize the bilayer within the ER (Gimenez-Andres *et al*., 2018).

#### (ii) The helix at the extreme C-terminal end of RBG multimers is amphipathic, except in SHIP164

At the other end of RBG multimers, all are immediately followed by a helix. In VPS13, ATG2, Tweek and most Hobbit homologs (not human, but in fly yeast and plant) the helices are amphipathic (Figure 6A, Supplementary Figure 4 and Supplementary Table 2B). These helices have biophysical properties similar to the helices that immediately precede the β- sheet (Figure 5), and they too are predicted by Alphafold to cross the RBG multimer and occlude it (for example, VPS13A see Figure 7), so they might have similar functions to their counterparts at the N-terminus. Future experiments will determine the extent to which these are to interact directly with the bilayer, for which feasible models have been created by others (Dall’Armellina *et al*., 2022), or alternately with protein partners as has been implied by functional relationships with scramblases (Ghanbarpour *et al*., 2021; Orii *et al*., 2021; Adlakha *et al*., 2022; Guillén-Samander *et al*., 2022).

**Figure 6.**
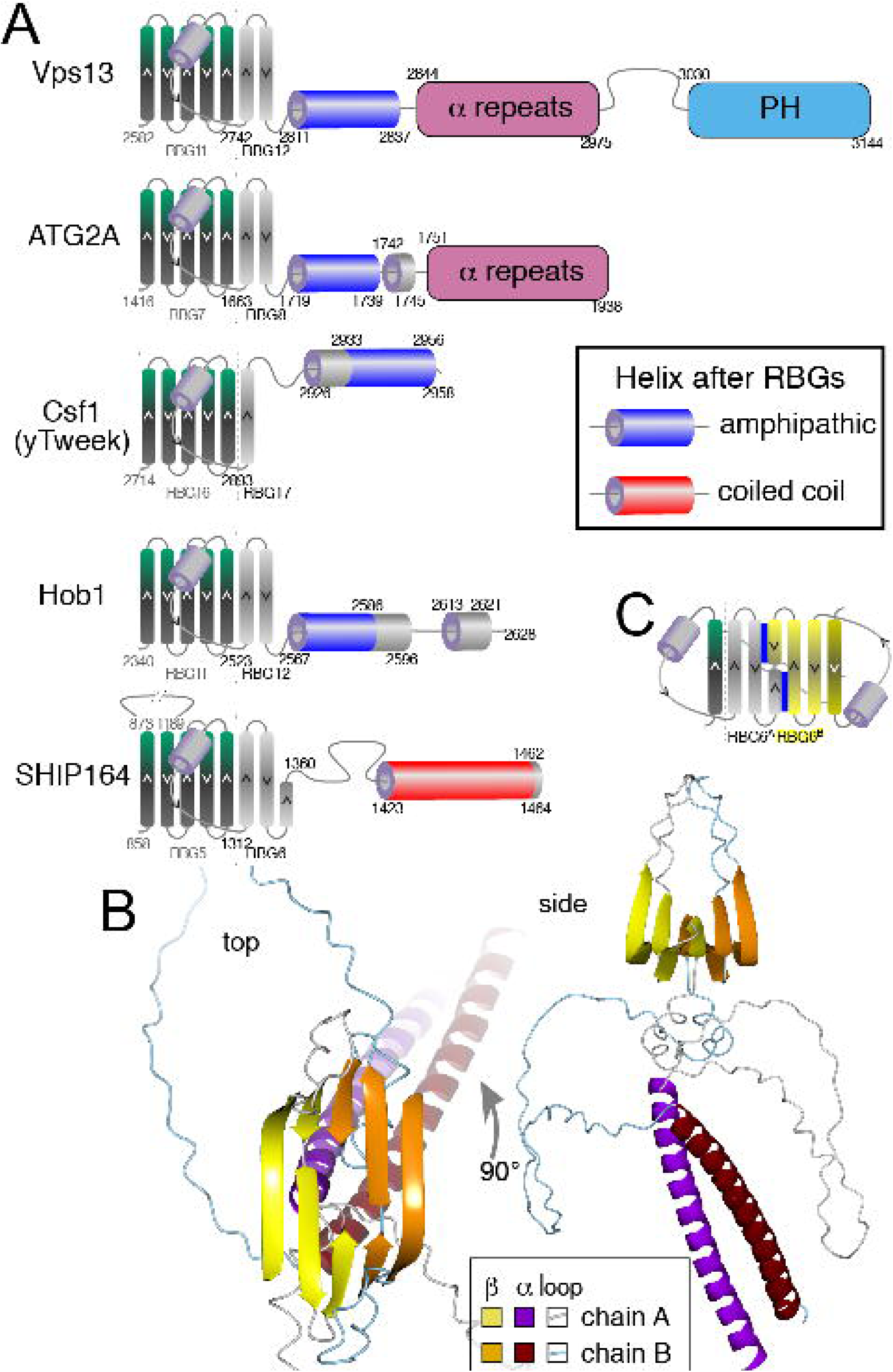
The C-terminus of the RBG multimer of SHIP164 uniquely has a coiled-coil. A. Helices following the final RBG domain in five families were assessed as amphipathic (blue) coiled coil (red) or neither (grey). For predicted helices be defined as amphipathic, 5 or more residues within a segment of 18 residues had to form a hydrophobic face (not glycine/alanine/proline). Recognized domains that follow the RBG multimer without any strongly predicted interaction are shown as filled shapes (not to scale, colors as Figure 1). The penultimate RBG domain is also shown for context. The examples chosen from each family are: VPS13: yeast; ATG2: human ATG2A; hobbit: yeast Hob1 (aka Fmp27); Tweek: yeast Csf1; SHIP164: human. Similar results were obtained with other family members. B. Top ranked model made by ColabFold of two copies of the C-terminus of SHIP164 (250 residues, 1215-1464), with RBG6 + terminal helix seen from the top (with fog as a depth cue) and from the side. Two dimerization interfaces were predicted, both with very high confidence (pLDDT≥90%): a β sheet (predicted for all 5 top models), and the coiled-coil (predicted in four of the five top models). Chains A and B are colored differently (see Key). C. Diagram of β-sheet dimeric interaction for SHIP164. The final strand of RBG5 and all of RBG6 are shown (colored as in A) together with a second copy rotated through 180° (colored yellow). The dimerization interface (blue lines) consists of parallel interactions between the final short strand and the N-terminal portion of the penultimate strand on the other monomer.

SHIP164 is the one RBG protein that has a different form. In opisthokont SHIP164 the linker after the final RBG ends with a coiled-coil. This means that the C-terminus of SHIP164 is unique among human RBG proteins by lacking a amphipathic helix.

#### (iii) Speculation about the C-terminus of SHIP164: might is dimerize end to end?

In addition to the C-terminus SHIP164 having a coiled-coil and amphipathic helix (Figure 6A), the RBG domain itself resembles a central domain, rather than a terminal one (Figure 3C). This suggests the possibility that C-terminus of RBG6 might have an alternate function to interacting with a membrane. Speculatively, it could interact with another RBG domain at a dimerization interface. ColabFold was therefore used to test whether SHIP164 can homodimerize. Predictions indicated two dimer interfaces: sheet-to-sheet and coiled-coil, (Figure 6B). The sheet interface is between the final strand, which is half-length and the penultimate strand (Figure 6C).

Although AlphaFold is imperfect, including in predicting dimeric interfaces (Bryant *et al*., 2022), these predictions taken as a whole are intriguing as they suggest the possibility that SHIP164 *in vivo* forms tail-to-tail dimers. Although such dimers have been reported for purified SHIP164 *in vitro* (Hanna *et al*., 2022), this observation is not conclusive because purified ATG2 also forms dimers at high concentrations. This question can only be addressed by seeking experimental evidence for the possibility of SHIP164 dimerization *in vivo*. In the absence of such evidence, it is worth pointing out that a plausible alternate possibility that also explains the bioinformatic findings is that SHIP164 is a monomer, interacting with a membrane partner, possibly via a hetero-dimeric coiled-coil. By contrast, homo-dimerization *in vivo* would have two functional implications: firstly, extending the reach of SHIP164 from ∼10 nm to ∼20 nm, consistent with the inter-membrane distances between endocytic vesicles enriched for SHIP164 (Hanna *et al*., 2022). Secondly, the two membrane interaction sites of a tail-to-tail dimer are identical copies of the N-terminus, which would imply symmetrical lipid transfer by SHIP164 bridging between homotypic organelles. Such homotypic activity has been described for cholesteryl ester transfer protein (CETP) acting on biochemically similar HDL particles (Mohammadpour and Akhlaghi, 2013). Here, homotypic activity by SHIP164 would involve bridges between the seemingly uniform endocytic vesicles that are separated from the ER by a zone of matrix (Hanna *et al*., 2022).

### D. Conservation at the level of sequence

#### (i) Amphipathic helices of the outermost RBG domains in VPS13 appear to have functions other than packing the terminal helices onto the groove

As described in Section B, there are four major types of RBG domain (RBG1/2/11/12, VPS13A numbering). These are also among the most highly conserved domains (Supplementary Figure 1). What are the unique structural and sequence features of these domains? To start with the outermost RBG domains (RBG1/12) were examined, focusing mainly on VPS13 as this family has the richest information. In both RBG1 and RBG12 the amphipathic helices cap the groove by crossing it perpendicularly to the sheet (red arrows in Figure 7A). This property of RBG1 has been seen by crystallography in VPS13 and ATG2 (Kumar *et al*., 2018; Osawa *et al*., 2019), but for RBG12 there is no crystallographic data, only AlphaFold prediction. Many of the most highly conserved residues in these domains are located on both the perpendicular helices and the adjacent sheet, indicating that they are involved in packing interactions. However, these are not the same as the residues that specify the amphipathic helices, since the hydrophobic face of the amphipathic helices is not involved in packing. Instead these residues, all moderately or highly conserved, do not point at the subjacent sheet and do not directly contact it (Figure 7B). This indicates that the amphipathicity of these helices is conserved as a separate feature. Speculatively, if the amphipathic helices do engage with a partner, either a lipid bilayer or a protein, they might adopt different conformations from the ones found in available structures. One possibility which might be explored is that they could rotate to open up the hydrophobic groove allowing lipid entry/exit (Figure 7B, curved arrows). Some support for helix mobility comes from considering Isoleucine 2771 in VPS13A, which causes disease when mutated (Figure 7B, grey arrow) (Rzepnikowska *et al*., 2017). The analogous mutation I2749R modelled in yVps13 reduces binding of the C-terminus to phosphatidylinositol 3-phosphate (PI3P). Given its location on the inner surface of strand 1 of VPS13A RBG12, the increased size and charge of the mutant side-chain could not affect PI3P binding directly, but might affect the closed position of the amphipathic helix.

#### (ii) Conserved structural variations differentiating between the two archetypal central RBG domains VPS13 vary across orthologous domains

The two RBG domain types of greatest significance are RBG2 and RBG11 (VPS13A numbering), because they represent archetypal forms in sequence terms that account for almost all central domains in VPS13, ATG2 and SHIP164 and for ∼50% of domains in Hobbit and Tweek (Section B, Figure 4). To understand these two major types of domain, the predicted structures RBG2 and RBG11 were compared (Figure 8A). This showed that each has a characteristic structural feature: RBG2 has a loop between strands 3 and 4 (≥15 aa), and RBG11 has a bulge inserted in the middle of strand 4 (≥5 aa) onto the external face of the groove (Figure 8A). The loop in RBG2 is verified by the identical appearance being seen by crystallography (Kumar *et al*., 2018). The bulge in RBG11 cannot be verified directly (but see below). Similar loops tend to be present in domains related by sequence to RBG2 and bulges in domains related to RBG11, including RBG6 where a bulge is present in the cryo-EM structure of (Li *et al*., 2020). Parallel structural variations (loop and the bulge) are smaller or absent in RBG domains otherwise homologous to RBG2 or RBG11, both in humans (Figure 8B) and in plants (Figure 3C, circles highlighting domain positions). Indeed, some domains have both features (Supplementary Figure 5A). Variability of the features is illustrated by RBG10, which has neither feature in any orthologous domain except for animal VPS13A/C, indicating that a loop may have been acquired just in that clade. Therefore, loops and bulges show too much variability to provide information about the events that created the domains present in LECA.

**Figure 7:**
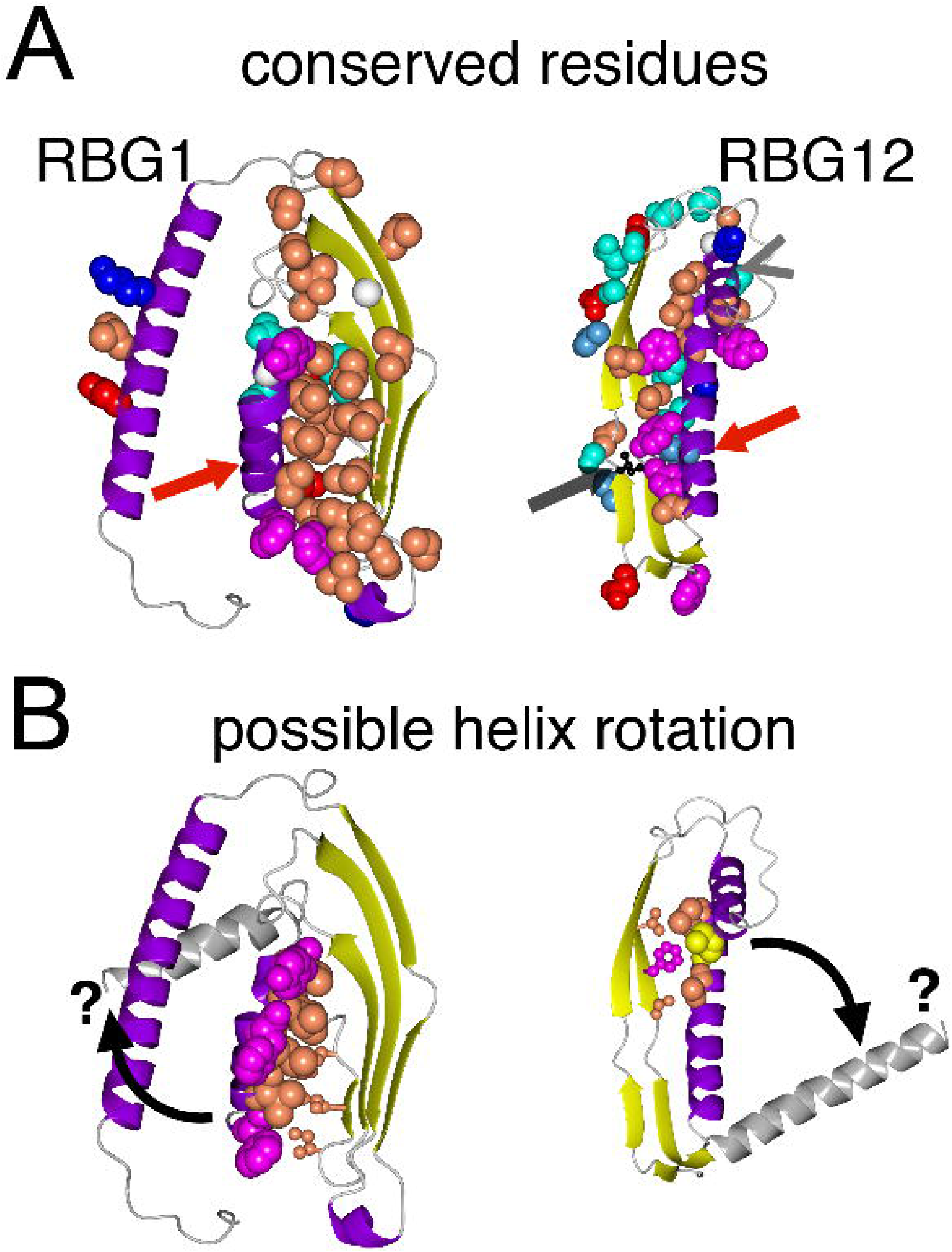
Conserved residues in VPS13 form a strip along the hydrophilic outside surface. A. and B: Models of RBG1 and RBG12 from VPS13, each domain rotated to highlight its interface between helix and sheet. **A.** shows the most conserved residues identified by Consurf are as spheres. Red arrows indicate the helices that run perpendicularly across the outermost ends of the RBG multimer (N-terminal helix in RBG1 and the C-terminal helix in RBG12). A helix is inserted between strands 1 and 2 of RBG12 (grey arrowheads in B). Color scheme:- secondary structure: sheet–yellow; helix–purple (partly transparent); loop– grey; side chains only of highly conserved residues (spheres) are colored by residue type: A/I/L/P/V: coral pink, FWY: magenta, NQST: cyan, G: white Cα, His: light blue, RK: blue, DE: red (using single letter code). Structures are based on AlphaFold models. **B.** highlights two small groups of residues: spheres = hydrophobic face of the amphipathic helices (RBG1: 1/3/7/8/11/14/15/18/19/22; RBG12 2852/6/9); ball and stick = sheet residues near to these (RBG1:27/29/31; RBG12 2821/3/5). There is no direct contact. Grey helices and black curved arrows and question marks indicate speculative movement of helices to open the ends of the groove. Isoleucine 2771, mutated to arginine in some patients, is indicated by the black ball and stick residue in A – see main text.

**Figure 8:**
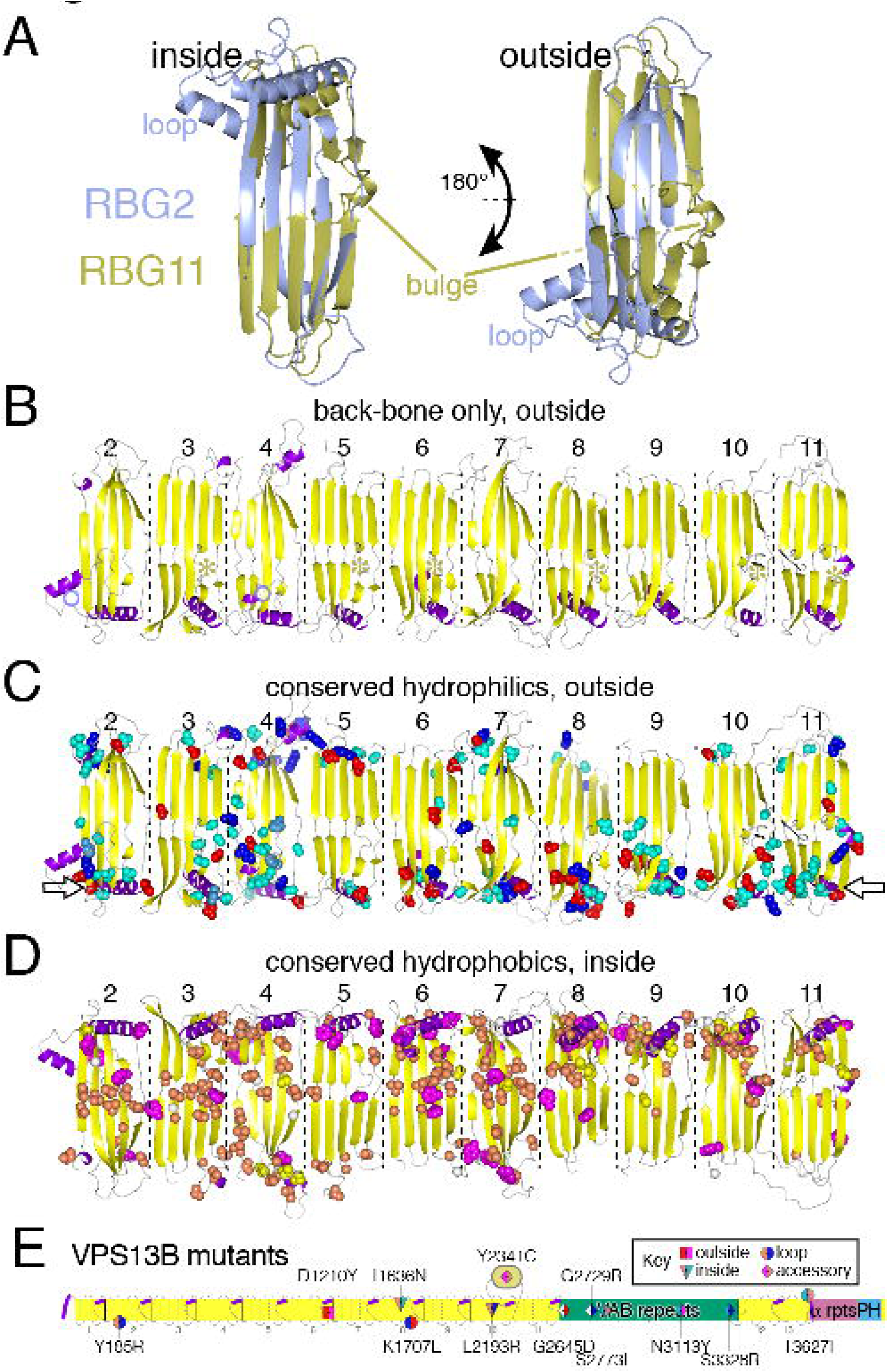
Distribution of conserved structural and sequence variations in VPS13 A: Comparison of the predicted structures of RBG2 (light blue) and RBG11 (gold) from VPS13A, indicating the extended loop between strands 3 and 4 of RBG2 and the bulge in strand 4 of RBG11; left hand view shows concave hydrophobic surface, right-hand view (rotated 180° over a horizontal axis), so both views have the N-terminus on the left-hand side. The loop in RBG2 includes a short helix (residues 228-236) that is placed adjacent to the outside surface of RBG1 both by AlphaFold and in the crystal structure 6CBC, however neither the helix nor the sheet has conserved residues. Black lines in RBG11 indicate positions of the start and end of the WWE domain (Figure 4A). B–D: Three different sets of views of the 10 central RBG domains from VPS13A (human, predicted in 2 segments, courtesy of Pietro De Camilli) aligned in the same orientation, untwisting the natural superhelix. The three views are: B. Back-bone only of convex surfaces (oriented as in right-hand view in A), showing loops as in RBG2 (light blue circles) and bulges as in RBG11 (gold asterisks); C. The most conserved hydrophilic residues (DEHKNQRST) showing only the concave surface (as in B); open arrows indicate the dominant location for these residues near the rim of the groove where the helices originate. D. The most conserved hydrophobic residues (ACFGILMPVWY) showing only the convex surface, view as in left-hand view in A). Spheres indicating conserved residues are colored as Figure 7. Structures are based on AlphaFold models, with the helical bunding (“handle”) removed for clarity. E. Locations of 12 VPS13B missense variants that cause human disease without effects on protein expression (Zorn *et al*., 2022). Location of mutated residue is indicated by shape as in the key: triangle (x2) = inside (hydrophobic) surface of groove; square (x1) = outside (hydrophilic) surface of groove; circle (x3): intrinsically disordered loop; diamond (x6) = accessory domain. Shapes are colored for wild-type (left) and mutant (residues) side-chains: ILC: coral pink; Y: magenta; NST: cyan; G: white; RK: blue; DE: red; numbering is for the 4022 aa VPS13B isoform, which has 25 aa inserted at ∼1400 aa compared to the isoform reported by Zorn *et al* (2022); domains are colored as Figure 1A, with added grey lines and thick purple segments to indicate loops and helices. Elements in domains (each RBG strand and each VAB repeat) are separated by dashed and dotted lines respectively. Note that the Y2341C mutation is in the β-sandwich inserted in the loop following RBG10 (Figure 4B).

#### (iii) Conserved residues in RBG domains VPS13 form a strip on the external, hydrophilic surface ideally placed for interacting with partners

As a final step to understanding RBG domain conservation, the most highly conserved residues in the central domains (RBG2–11) of VPS13A were mapped (Figure 8C and D, and Supplementary Movies 1–3). Looking first at the helices at the ends of domains, half of RBG2–11 have conserved residues packing the interface between the helix and adjacent sheet, while the other half have no or few conserved residues on the helix or sheet (Supplementary Figure 5D). This suggests that such helices might be mobile, however, countering this, in cryo-EM of VPS13 all the helices of RBG domains 2 to 7 align with the long axis contacting the rim of the groove (Li *et al*., 2020), which suggests that the helices are not mobile.

A second feature of the central domains is the presence of conserved residues along the entire outside convex surface of the groove, which is simplified here by showing hydrophilic residues only on the convex surface and hydrophobic residues only on the concave surface (Figure 8C and D), and is also seen for all residues on all surfaces (Supplementary Movies 2 and 3). To check if conserved residues near the C-terminus are involved in intramolecular interactions with accessory domains, RBG10/11/12 were examined for contacts with VAB (repeat 1) ATG_C and PH domains, which AlphaFold places overlying them. Analysis of co-evolution maps indicated that there is no contact (data not shown).

Similar to VPS13, conserved residues on external faces are found not only for central domains of other RBG multimers (Supplementary Figure 5A–C), but also across almost all of Hobbit (Supplementary Figure 5D and E). Overall, these findings indicate that a large majority of the entire length of RBG multimers has conserved residues on their outside (mainly hydrophobic) surfaces. Conserved hydrophilic residues on the convex surfaces of RBG domains tend to occur near the rim of the groove where helices originate (Figure 8C, open arrows), which suggests that this rim is a characteristic site for interactions with partners. With the exception of one-two RBG domains per protein, this applies to the most central RBG domains, which are furthest from membrane anchoring points. An example of the importance of such central conservation is found in a catalog of high probability disease-causing nonsense mutations in VPS13B (Zorn *et al*., 2022). While 2 out of 12 mutated sites are in hydrophobic residues that line the groove, 4 are external facing residues located along various parts of the multimer and six in accessory domains, particularly the VAB repeats (Figure 8E). Thus, disease genetics agrees with sequence conservation to suggest that residues far from membrane anchoring points that do not interact with lipid cargo are critical to function.

## Conclusions

Analyzing the sequences of RBG protein sequences has led to the examination of several conserved generic properties: the number of RBG domains; patterns of internal duplication of domains; accessory domains; interaction modules at the extreme ends of the multimers; and conserved residues along the entire RBG multimer. Because all of these properties are conserved, they are likely to be related to conserved functions. The number of domains may match the width of contact site, which will surely be a subject of future study, along with multimer flexibility to accommodate changes in length. The pattern of domain duplication shows however much of the origins of the RBG superfamily as can still be detected in sequences >1.2Bn years after they originated (Mills *et al*., 2022). One major caveat to these conclusions is that they assume that current sequences indicate a pattern of gradual accumulation of domains from ancestral forms related to RBG1-2-11-12 vertically across time, rather than events such as gene conversion either within or between RBG proteins horizontally. However, whatever the events were, conservation across eukaryotes places them as taking place before the formation of LECA.

As domains duplicated, their binding partners might also have duplicated; for example multiple members of protein family X may bind to different parts of VPS13 if the ancestral protein X interacted with either the ancestral form of RBG2 in RBG11. Folded accessory domains are strong candidates to mediate partner interactions, after the identification of key interactions for VAB repeats (Bean *et al*., 2018; Dziurdzik *et al*., 2020; Adlakha *et al*., 2022) and for the C-terminal PH domain of VPS13A (Guillén-Samander *et al*., 2022). This work defines the full range of such domains: the ones with defined folds are all in VPS13, mostly near its C-terminus. Amphipathic helices are present at the extreme ends of multimers, with one exception: the C-terminus of SHIP164, which instead has a coiled coil, the significance of which is not yet known. Sequence conservation on the external face of the lipid binding hydrophobic groove may participate in a series of binding sites along the entire bridge for protein partners. Some partners of RBG may be proteins that are separately recruited to the contact site, but others may be recruited solely by this interaction, potentially making RBG proteins hubs for contact site function.

Shortcomings of the study include that it only focusses on folded domains, leaving out short linear motifs, only some of which have been mapped in VPS13 (Kumar *et al*., 2018; Guillen-Samander *et al*., 2021) and ATG2 (Bozic *et al*., 2020; Ren *et al*., 2020), which fits with their overall discovery rate of <5% (Davey *et al*., 2017). Adding these to information on domain structure will allow further hypotheses on RBG protein function to be generated and tested.

## Methods

### Structural predictions - ColabFold

For predictions spanning RBG 10 and 11 of VPS13, the yeast Vps13p (yVps13) sequence from 1643-2840 was submitted to ColabFold after removing 2112-2542 (767 aa remaining) (Mirdita *et al*., 2021). This region aligns with human VPS13A 1635-2119+2502-2865. The full VPS13 model in Figure 2D contains: VPS13A (rat, 1-1642), yVps13 1657-2835 (residues 15-737 of the ColabFold model above) and residues 2836-3144 from a model of yeast Vps13 2580-3144 made by trRosetta (Yang *et al*., 2020). These three segments were aligned from overlapping segments in QtMG for Mac (CCP4MG v. 2.10.11). Models of other regions, including specific RBG domains, homodimers of C-terminal RBG domains plus following helices, and accessory domains were made in ColabFold as described in the text.

### Remote homology: HHpred

Sequences were obtained from both Uniprot and NCBI Protein databases for VPS13 in 8 organisms: human, fly (*D. melanogaster*), nematode worm (*C. elegans*), *Trichoplax adherens*, *S. cerevisiae*, *S. pombe*, *Capsaspora owczarzaki* and *A thaliana*. For the one sequence that was incomplete, *Trichoplax* VPS13A/C (1299 aa), additional sequence was assembled from adjacent genes that encode sequences homologous to the N- and C-termini of VPS13AC. This added the N-terminus, but left a likely gap of 1500-2000 amino acids that is possibly be encoded in a genomic region of 13891 bp. tBLASTn in this region identified 262 aa of VPS13A/C-like sequence in 6 regions (probable exons, data not shown) in 5 statistically significant hits, suggesting that an unannotated complete VPS13A/C is expressed in *Trichoplax*, of which 1561 aa have so far been identified.

The form that RBG domains take in HHpred (Soding, 2005) was initially determined from cross-correlating HHpred searches seeded with portions of entire protein structures predicted by AlphaFold. Many strands are seen in HHpred as two disconnected halves, each 6-10 residues. Inserts with no sheet or helix were ignored. Long inserts in the middle of domains were omitted if they prevented alignment on one side of the insert. This allowed identification of all RBG domains in human VPS13A, VPS13B, extra RBG domains in VPS13C/D, and selected domains from yeast and *Arabidopsis.* Benchmarking of HHpred against AlphaFold showed that HHpred underestimates the length of β-strands in RBG domains by >10%, and that it mis-identifies a small minority of strands as helices (data not shown). Based on these observations, all HHpred hits longer than 25 residues with predicted shared structure ≥10% that contained any predicted sheet were considered true positives.

Each domain, defined as the strands plus 12-20 aa of the 6^th^ element (*i.e.* the first few turns of the helix for uniformity), was submitted to HHpred with 5 iterations of HHblits to make multiple sequence alignments (MSAs), then used to find homologs in target HMM libraries of key whole proteomes (human, *S. cerevisiae*, *C. owczarzaki* and *A thaliana)*. To identify homology of a query domain to RBG2 and RBG11 in VPS13, four pair-wise alignments were made using the “Align two sequences/MSAs” option, aligning the query with MSAs from the penultimate domains both of VPS13A (RBG2 &11) and of VPS13B (RBG2 & 12). Both VPS13A and VPS13B were included here to capture the breadth of VPS13 sequences, and the four MSAs of penultimate domains were made superficially (one iteration of HHblits) to preserve unique qualities of the domains.

All regions that did not align well with the typical 5 strand+loop pattern of RBG domains were submitted to HHpred to identify their fold (Gabler *et al*., 2020), and also had structure predicted by ColabFold (Mirdita *et al*., 2021).

### Cluster Maps

For relationships between RBG domains, 229 domains from VPS13 proteins in the 8 eukaryotes listed above were submitted to CLANS, using BLOSUM45 as scoring matrix (Frickey and Lupas, 2004; Gabler *et al*., 2020). 162 connected domains at p<0.1 were clustered (>100,000 rounds) in 2 dimensions.

For relationships between whole VPS13 proteins, 1266 full length proteins sequences were submitted to CLANS, using BLOSUM62. This list was obtained by generating two separate lists of proteins homologous to the central portion of the RBG multimer (to avoid proteins that only are homologous to the VAB domain) seeding HHblits searches with either VPS13A/C/D (RBG2-9) or VPS13B (RBG2-10) (Remmert *et al*., 2012). The resulting lists of 886 VPS13A/C/D homologs (1 iteration) and 703 VPS13B homologs (3 iterations) were then reduced by removing (near) identities using MMseq2 with default settings (Steinegger and Soding, 2017), and the four human sequences were added. Full-length sequences were used to cluster, >100,000 rounds, with 1186 proteins connected at p<10^-30^.

### Motif Consensus

Diverse sequences at N-terminus of ATG2 were gathered using HHblits, and the N-termini (39 columns) were aligned with MUSCLE (www.ebi.ac.uk), and visualized with WebLogo (weblogo.berkeley.edu).

### Analysis of Helices and short sequences

Sequences were analyzed for propensity to be amphipathic and coiled coils using Heliquest and Coils tools, respectively (Lupas *et al*., 1991; Gautier *et al*., 2008). A consensus from the N-termini of ATG2 homologs was obtained from a MSA after 3 rounds of HHblits seeded with ATG2 from *Drosophila* (Remmert *et al*., 2012). From 514 entries, sequences containing intact extreme N-termini (38 aa) were selected, rare inserts were reduced by dropping sequences plus editing by hand, and identical sequences reduced to single entries, leaving 46 sequences. The first 9 columns of this were submitted to WebLogo (https://weblogo.berkeley.edu).

### Identification of Conserved Residues

AlphaFold models, either complete or divided into regions defined as RBG domains (see Supplementary File 1) were submitted to Consurf (Ashkenazy *et al*., 2016), using standard settings except breadth of searches was maximized by setting HMMER to 5 iterations, producing 400-2000 unique sequence homologs. Where insufficient homologs were obtained, HHblits was used to build an MSA – either 2 or 3 iterations, n=200-300. Highly conserved residues were identified as those scoring 8 or 9 on conservation color scale (from 1 to 9).

## Supplementary Information: 5 Figures, 2 Tables, 1 Data File and 3 Movies

### Supplementary Figures

Supplementary Figure 1. RBG domain cluster map identifies distance between duplicates of RBG5 and RBG6.

Cluster map made in CLANS of 162 linked RBG domains from VPS13 proteins from 7 diverse eukaryotes (see Methods). Stronger homologies, shown by short and thick lines, is confined to orthologous domains. All RBG7s clustered tightly, including RBG7C/D^1/2^. By comparison, RBG5C^1/2^ and RBG6C^1/2^ form two clustered groups separate from RBG5D^1/2^ and RBG6D^1/2^ respectively, and in both RBG5 and RBG6 D^1^ is only indirectly or weakly connected with D^2^. This suggests that the duplication in VPS13D was a separate earlier event from that in VPS13C.

Supplementary Figure 2. VPS13B homologs are spread across eukaryotic evolution

Cluster map of 1186 VPS13 proteins constructed in CLANS as described in Methods. VPS13B and close homologs clustered separately from VPS13A/C/D, in which VPS13D form one segment. The VPS13B cluster contains 25 protist proteins (blue): Albugo, A0A024G1K2; Aphanomyces, A0A397D2M9, A0A418DCD9, A0A397ERH8, A0A024TE03, A0A6A4ZLW7; Bremia, A0A484ECX3; Chara, A0A388LFT4; Chromera, A0A0G4FFP3; Globisporangium, K3X2H6; Guillardia, A0A7S4JEF6; Hanusia, A0A7S0E6H4; Hemiselmis, A0A7S0TJS0; Hondaea, A0A2R5GB42; Hyaloperonospora, M4BW50; Klebsormidium, A0A1Y1HLB4; labyrinthulid, A0A7S2S870; Phytophthora, A0A0W8B040, A0A081AUX5, A0A0W8D6K5, H3H1A7, A0A3F2RKP7; Pythium, A0A2D4C9P6; Thecamonas, A0A0L0DHF4; Vitrella, A0A0G4GLM3.

Supplementary Figure 3. RBG domain 5 of VPS13X from *A. thaliana* has 7 strands.

Residues 939-1181 of VPS13X from *A. thaliana* At3g50380 were modelled by ColabFold, and visualized both in ribbon form showing the inside of the groove (top, strands – yellow, helices – purple) and with the surface of the sheet colored according to the YRB scheme (see key (Hagemans *et al*., 2015)) indicating its hydrophobicity, together with two other orientations showing the side and hydrophilic outside of the groove. pLDDT of the model (excluding inserted loops) was 83% over 174 residues.

Supplementary Figure 4. Amphipathic helices at the extreme C-termini of RBG multimers

A–D Helical wheel projections (from Heliquest) of the indicated residues immediately following the final RBG domain of VSP13A (human), ATG2A (human), yeast Tweek (Csf1) and fly Hobbit respectively. Images created by Heliquest, with hydrophobic face indicated by an arc.

Supplementary Figure 5. Conserved residues on the concave (outside) face of domains near the central regions of RBG proteins

A–C: Concave (outside) faces of: A. ATG2 RBG5, B. SHIP164 RBG4, C. yeast Tweek (Csf1) RBG12. Identification and coloring of conserved residues (all), colored as in Figure 7. ATG2 RBG5 has both a loop between strands 3 and 4 (light blue arrow) and a bulge onto the convex surface between the two halves of strand 4 (gold arrow). D. 10 central RBG domains from VPS13A rotated to view helices and adjacent sheet, with all conserved residues shown as spheres, coloring as in Figure 7, same models as Figure 8B–D. Note that conserved residues pack the interface between helix and sheet in half of the RBG domains: RBG3/6/8/9/10. E and F: Yeast Hob1 (aka Fmp27) showing conserved hydrophilic (E) and hydrophobic (F) residues, as in Figure 8C/D, with views rotated by 180° in both cases. Three pairs of antiparallel helices have been removed. Open arrows indicate a region with few conserved residues 559-892 (strands 4/5 of RBG4 and all of RBG5). This differs from the orthologous region of human Hobbit, where RBG5 shows wide homology, for example to its plant ortholog (Figure 3E).

Supplementary Table 1: Homology of RBG domains in VPS13A/B, ATG2B and SHIP164 to the penultimate domains in VPS13 (RBG2 at the N-terminus and RBG11/12 at the C-terminus of VPS13A/B).

RBG domains in the indicated proteins were compared pairwise in HHpred with penultimate domains of VPS13A and VPS13B, as explained in the Methods. pSS >90% = hit (strong); 40-90% = moderate; 10-40% = weak; ≤10% = no hit. “/” indicates no hit or probability shared structure (pSS)<1%. 1st and last RBG domains are included only for VPS13A. * inserts were removed from VPS13A RBG11 (WWE domain) and SHIP164 RBG5 (disordered loop). VPS13C/D follow VPS13A (except for the inserted 3 domains).

Supplementary Table 2: Amphipathic helices immediately before and after the RBG multimer Helices predicted by HHpred and/or AlphaFold, not overlapping with TMHs identified by TOPCONS, were tested in Heliquest typically using an 18-mer window, reduced in width when necessary.

Supplementary Data File 1. Domain boundaries for RBG proteins in this study

Boundaries of all RBG domains and non-standard accessory domains in selected proteins for this study. The C-terminal boundaries include the initial portion only (12-20 aa) of the 6^th^ element (loop).

## Movies

RBG domains only from human VPS13A (N-terminus to the left) showing: Movie 1 back bone only, Movie 2 conserved hydrophilic residues, Movie 3 hydrophobic residues.

## Supplementary Materials

Underlying research materials that support this study, in particular the original HHpred searches, will be made freely available in Harvard Dataverse at https://dataverse.harvard.edu/dataverse/RBG_Superfamily

## Acknowledgments

Thanks to Rosario Valentini and Pietro De Camilli for sharing PDB files prior to publication respectively of complete Csf1 and human VPS13A. I would like to thank Matt Hayes, Arash Bashirullah and Sarah Neuman for discussions about the manuscript.

