## Supplementary figures and images for "Sequence analysis and structural predictions of lipid transfer bridges in the repeating beta groove (RBG) superfamily reveals past and present domain variations affecting form, function and interactions of VPS13, ATG2, SHIP164, Hobbit and Tweek"

### Supplementary Figure 1

#### Supplementary Figure 1

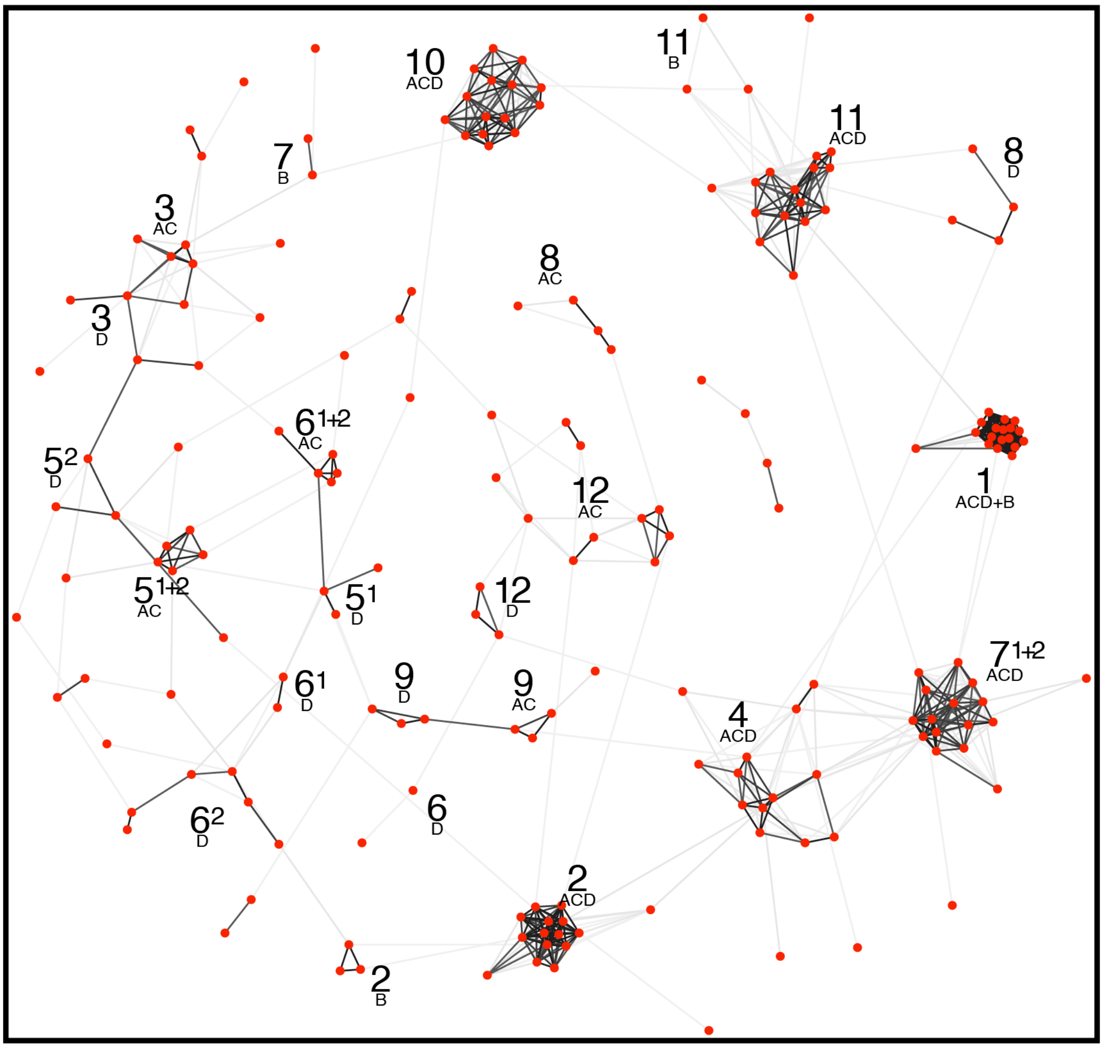

### Supplementary Figure 2

#### Supplementary Figure 2

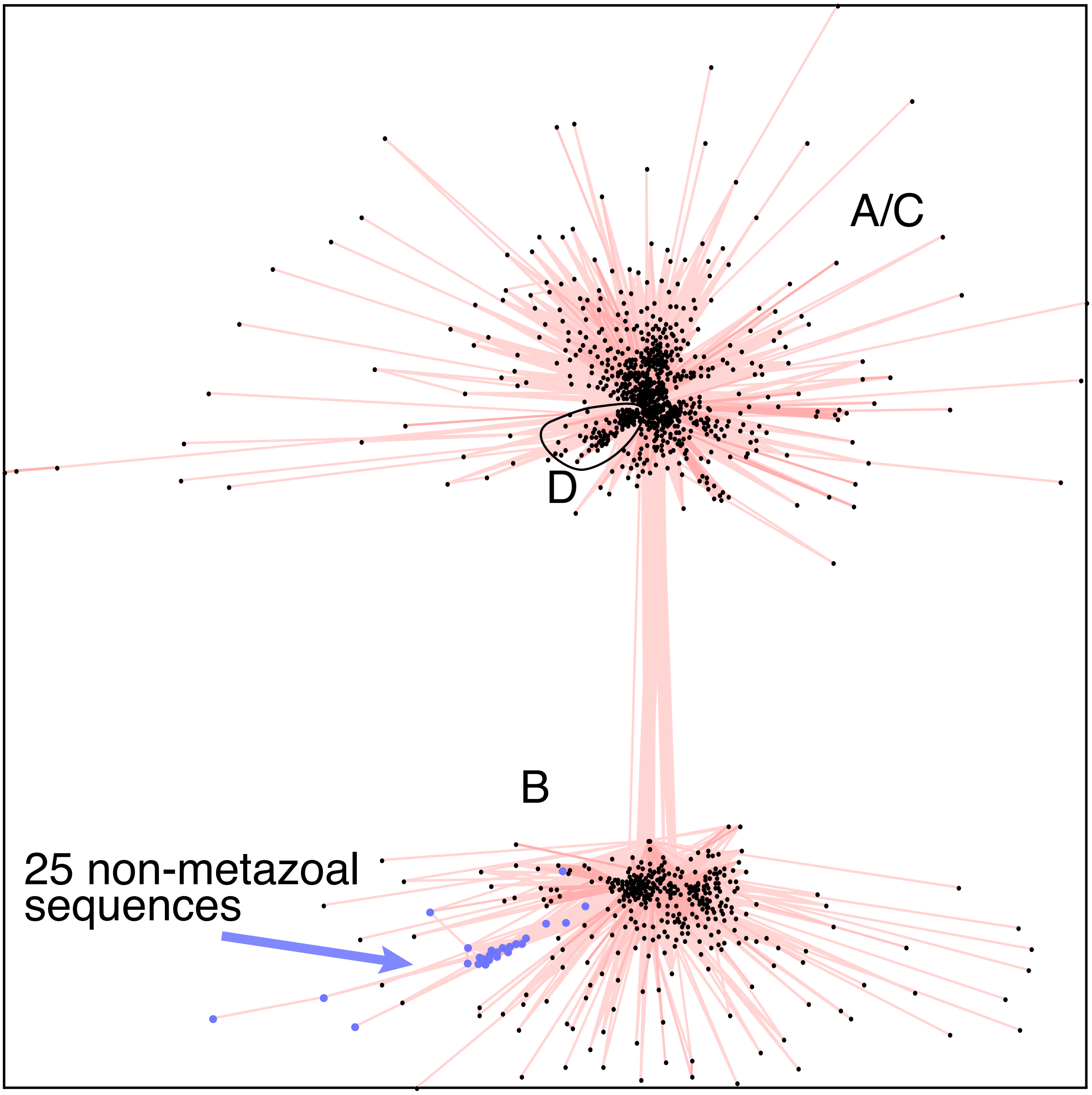

### Supplementary Figure 3

#### Supplementary Figure 3

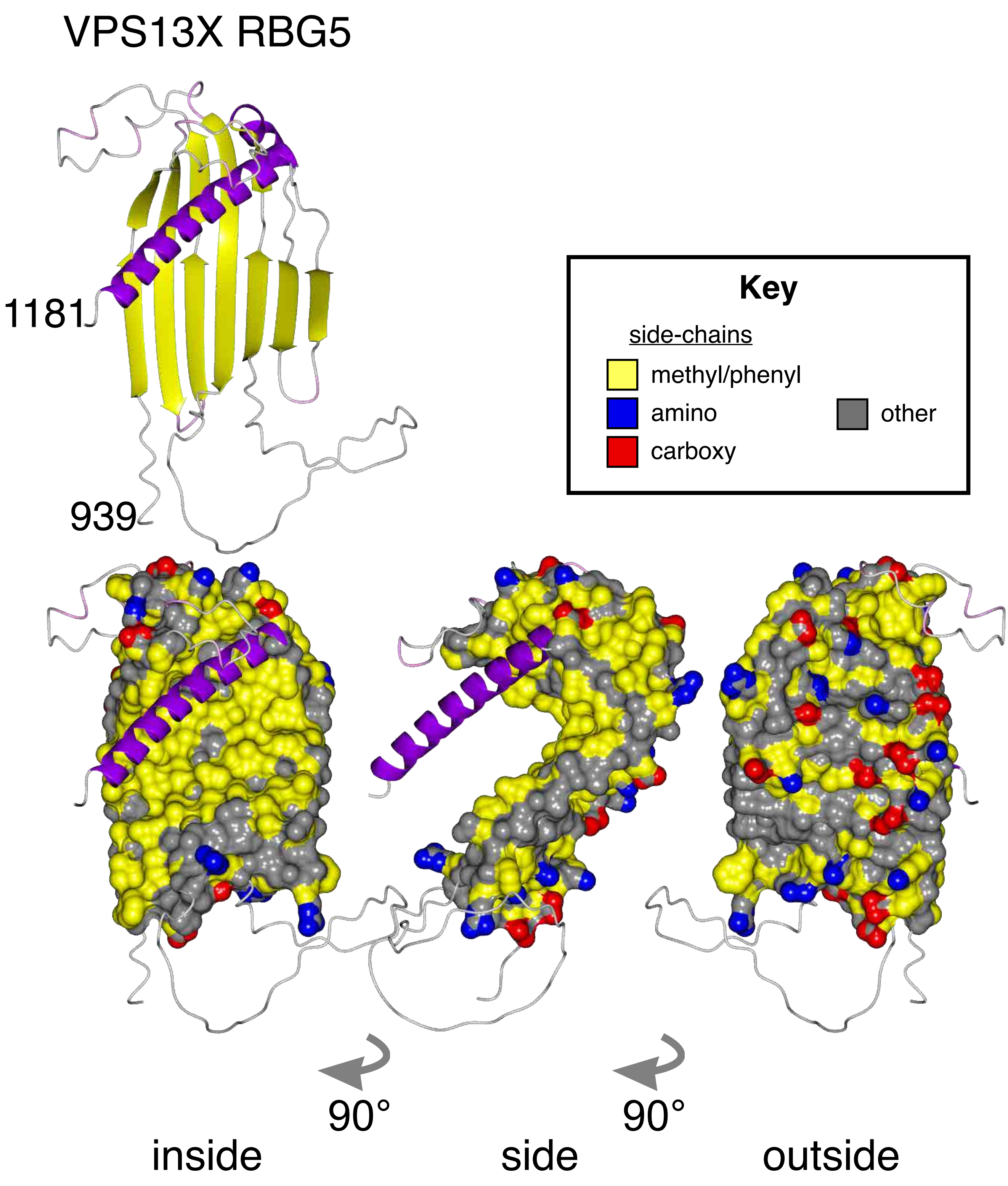

### Supplementary Figure 4

#### Supplementary Figure 4

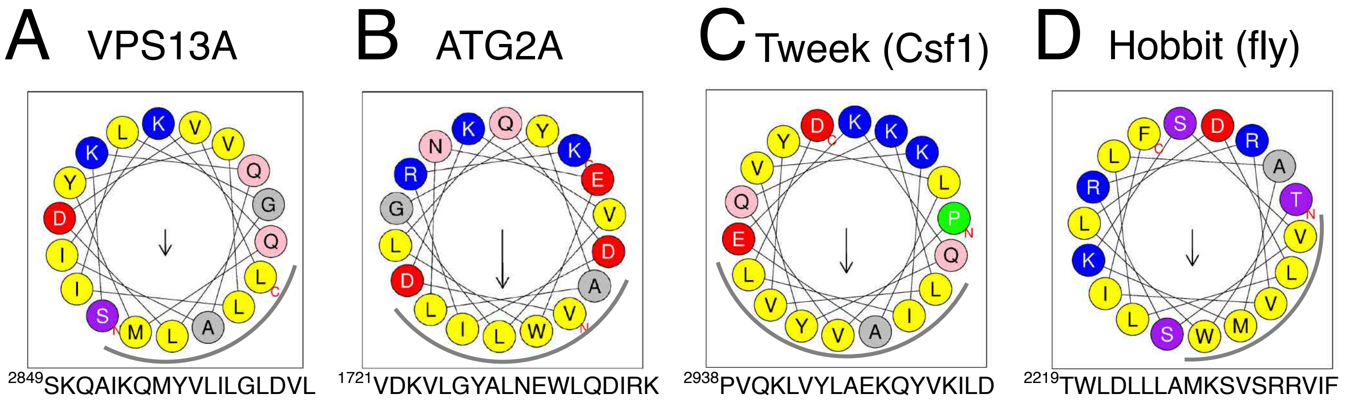

### Supplementary Figure 5

#### Supplementary Figure 5

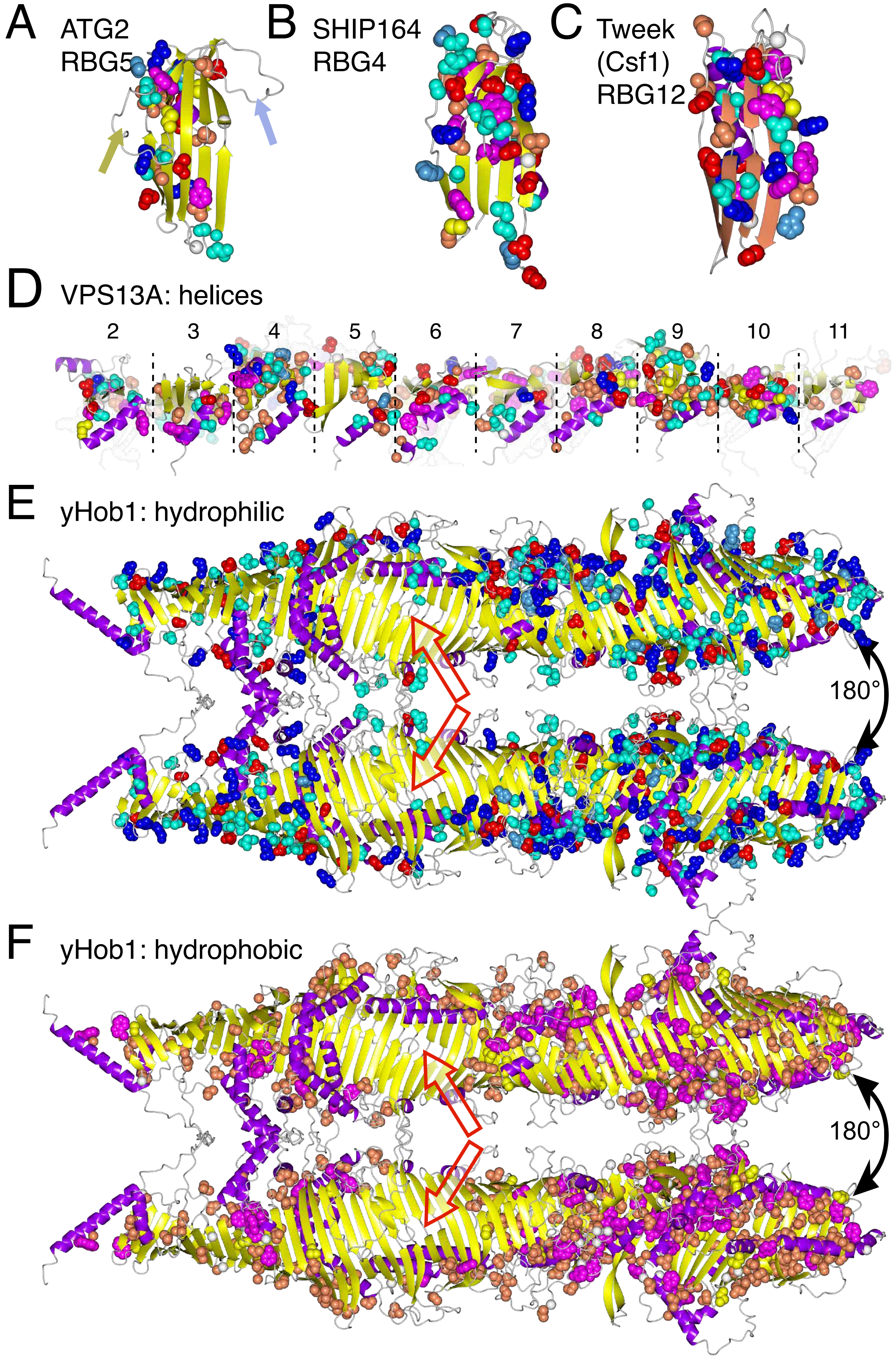
