## Supplementary Table 1 for "Sequence analysis and structural predictions of lipid transfer bridges in the repeating beta groove (RBG) superfamily reveals past and present domain variations affecting form, function and interactions of VPS13, ATG2, SHIP164, Hobbit and Tweek"

**Supplementary Table 1:**
Homology of RBG domains in VPS13A/B, ATG2B and SHIP164 to the penultimate domains in VPS13 (RBG2 at the N-terminus and RBG11/12 at the C-terminus of VPS13A/B).

RBG domains in the indicated proteins were compared pairwise in HHpred with penultimate domains of VPS13A and VPS13B, as explained in the Methods. pSS >90% = hit (strong); 40-90% = moderate; 10-40% = weak; ≤10% = no hit. "/" indicates no hit or probability shared structure (pSS)<1%. 1st and last RBG domains are included only for VPS13A. * inserts were removed from VPS13A RBG11 (WWE domain) and SHIP164 RBG5 (disordered loop). VPS13C/D follow VPS13A (except for the inserted 3 domains).

A

| VPS13 | | RBG 2 = N-term. (VPS13_N domain) | | | | RBG 11/12 = C-term. (part of Apt1 domain) | | | | notes |
|---|---|---|---|---|---|---|---|---|---|---|
| | | VPS13A | | VPS13B | | VPS13A* | | VPS13B | | |
| | RBG | pSS | cols | pSS | cols | pSS | cols | pSS | cols | |
| A | 1 | 5 | 15 | 7 | 13 | / | | / | | Chorein_N (100%) |
| | 2 | 100 | 156 | 100 | 126 | 6 | 24 | 1 | 24 | RBG2 |
| | 3 | 7 | 38 | 1 | 34 | 99 | 115 | 93 | 68 | RBG11/12 |
| | 4 | 97 | 73 | 88 | 53 | / | | / | 23 | RBG2 |
| | 5 | 2 | 44 | / | | 96 | 120 | 21 | 65 | RBG11/12 |
| | 6 | 2 | 17 | / | | 98 | 115 | 38 | 54 | RBG11/12 |
| | 7 | 98 | 126 | 98 | 107 | / | | / | | RBG2 |
| | 8 | 1 | 15 | 4 | 38 | 98 | 114 | 97 | 122 | RBG11/12 |
| | 9 | 0 | 111 | 40 | 65 | 8 | 111 | 18 | 57 | RBG2 (mixed) |
| | 10 | 66 | 25 | 77 | 112 | 9 | 27 | 4 | 25 | moderate RBG2 |
| | 11* | 3 | 19 | 4 | 28 | 100 | 123 | 100 | 123 | RBG11/12 |
| | 12 | 2 | 7 | / | | / | | 2 | | VPS13_C |
| B | 2 | 100 | 126 | 100 | 128 | 3 | 22 | 9 | 23 | RBG2 |
| | 3 | / | | 1 | 50 | 8 | 21 | 15 | 43 | weak RBG11/12 |
| | 4 | 26 | 51 | 85 | 81 | / | | / | | moderate RBG2 |
| | 5 | 35 | 35 | 47 | 108 | 83 | 106 | 81 | 117 | mod. RBG11/12 (mixed) |
| | 6 | 93 | 53 | 96 | 104 | 1 | 17 | / | | RBG2 |
| | 7 | 2 | 27 | 2 | 35 | 83 | 127 | 80 | 146 | moderate RBG11/12 |
| | 8 | 3 | 19 | 5 | 72 | 2 | 48 | 2 | 38 | none |
| | 9 | 1 | 16 | 1 | 77 | 92 | 102 | 94 | 103 | RBG11/12 |
| | 10 | 79 | 56 | 97 | 73 | 2 | 26 | 2 | 76 | RBG2 |
| | 11 | 70 | 51 | 74 | 73 | / | | 4 | 43 | moderate RBG2 |
| | 12 | 10 | 25 | 7 | 25 | 100 | 127 | 100 | 163 | RBG11 |
| X | 2 | 91 | 115 | 96 | 114 | 3 | 19 | 6 | 53 | RBG2 |
| | 3 | / | | 1 | 16 | 28 | 55 | 27 | 62 | weak RBG11/12 |
| | 4 | 16 | 18 | 40 | 35 | 1 | 16 | 1 | 6 | weak RBG2 |
| | 5 | / | | / | | 1 | 15 | / | | none |
| | 6 | 20 | 53 | 9 | 52 | 2 | 15 | 2 | 15 | weak RBG2 |
| | 7 | 1 | 16 | 20 | 43 | 37 | 109 | 31 | 39 | weak RBG11/12 (mixed) |
| | 8 | 6 | 23 | 2 | 22 | 99 | 123 | 99 | 125 | VPS13_C |

B

| ATG2B | RBG 2 = N-term. (VPS13_N domain) | | | | RBG 11/12 = C-term. (part of Apt1 domain) | | | | notes |
|---|---|---|---|---|---|---|---|---|---|
| | VPS13A | | VPS13B | | VPS13A* | | VPS13B | | |
| RBG | pSS | cols | pSS | cols | pSS | cols | pSS | cols | |
| 1 | 14 | 16 | 17 | 15 | / | | 1 | 6 | Chorein_N (100%) |
| 2 | 98 | 152 | 99 | 126 | / | | / | | RBG2 |
| 3 | 4 | 18 | 1 | 20 | 16 | 83 | 22 | 46 | weak RBG11/12 |
| 4 | / | | 9 | 26 | 4 | 41 | 2 | 22 | none (and variable) |
| 5 | / | | / | | / | | / | | none |
| 6 | 34 | 118 | 67 | 116 | 1 | 14 | 3 | 9 | moderate RBG2 |
| 7 | 10 | 28 | 15 | 28 | 97 | 106 | 97 | 119 | RBG11 |
| 8 | 1 | | 1 | | 2 | 8 | / | | =V13B RBG13 p94% (68 cols) |

C

| SHIP164 | RBG 2 = N-term. (VPS13_N domain) | | | | RBG 11/12 = C-term. (part of Apt1 domain) | | | | notes |
|---|---|---|---|---|---|---|---|---|---|
| | VPS13A | | VPS13B | | VPS13A | | VPS13B | | |
| RBG | pSS | cols | pSS | cols | pSS | cols | pSS | cols | |
| 1 | 13 | 15 | 19 | 15 | / | | / | | Chorein_N (100%) |
| 2 | 97 | 126 | 99 | 125 | / | | 1 | 21 | RBG2 |
| 3 | 1 | 26 | 16 | 79 | 9 | 46 | 3 | 42 | ATG2 RBG7 |
| 4 | 58 | 54 | 80 | 55 | / | | / | | moderate RBG2 |
| 5* | / | | / | | / | | / | | ATG2 RBG7 |
| 6 | 68 | 39 | 87 | 34 | 4 | 14 | 1 | 14 | moderate RBG2 |
