## Supplementary Table 2 for "Sequence analysis and structural predictions of lipid transfer bridges in the repeating beta groove (RBG) superfamily reveals past and present domain variations affecting form, function and interactions of VPS13, ATG2, SHIP164, Hobbit and Tweek"

**Supplementary Table 2:**
Amphipathic helices immeidately before and after the RBG multimer

Helices predicted by HHpred and/or AlphaFold, not overlapping with TMHs identified by TOPCONS, were tested in Heliquest typically using an 18 mer window, reduced in width when necessary.

A. N-termini

| | | length | start | end | sample helix | |
|---|---|---|---|---|---|---|
| | | | | | hydrophobic face | charge |
| VPS13 | yeast | 21 | 4 | 24 | 8/18 | 0 |
| | human A | 21 | 5 | 25 | 8/18 | -2 |
| | human B | 20 | 4 | 23 | 9/18 | +3 |
| | human C | 21 | 5 | 25 | 8/18 | -2 |
| | human D | 21 | 4 | 24 | 8/18 | +1 |
| | A. thaliana S | 25 | 3 | 27 | 9/18 | 0 |
| | A. thaliana M1 | 22 | 4 | 25 | 10/18 | +1 |
| | A. thaliana M2 | 22 | 4 | 25 | 10/18 | +1 |
| | A. thaliana X | 19 | 4 | 22 | 5/18 | +6 |
| ATG2 | yeast Atg2 | 13 | 7 | 19 | 4 / 11 | +2 |
| | human A | 11 | 12 | 22 | 4 / 11 | +2 |
| | human B | 13 | 7 | 19 | 4 / 11 | +3 |
| | A. thaliana | 13 | 9 | 21 | 5 / 11 | +1 |
| SHIP164 | human | 22 | 2 | 23 | 5 / 18 | +5 |
| | Dictyostelium | 21 | 2 | 22 | 6 / 18 | +6 |
| | A. thaliana | 22 | 2 | 23 | 7 / 18 | +2 |
| Tweek | yeast | 25 | 36 | 60 | 6 / 18 | +1 |
| | human | 16 | 50 | 65 | 7 / 16 | +3 |
| | Trichomonas | 16 | 35 | 50 | 6 / 16 | +2 |
| Hobbit | yeast Fmp27 | / | | | | |
| | yeast Hob2 | / | | | | |
| | human | / | | | | |
| | A. thaliana | / | | | | |

(see over for part B)

Supplementary Table 2 continued:

B. C-termini

| | | length | start | end | sample helix | |
|---|---|---|---|---|---|---|
| | | | | | hydrophobic face | charge |
| VPS13 | yeast | 25 | 2812 | 2836 | 5/18 | 0 |
| | human A | 28 | 2842 | 2869 | 5/18 | +1.5 |
| | human B | 25 | 3628 | 3652 | 8/18 | +1 |
| | human C | 24 | 3396 | 3419 | 5/18 | +1 |
| | human D | 23 | 4044 | 4066 | 5/18 | -1 |
| | A. thaliana S | 20 | 3088 | 3107 | 5/18 | +1 |
| | A. thaliana M1 | 23 | 3775 | 3797 | 5/18 | +2 |
| | A. thaliana M2 | 23 | 3648 | 3670 | 8/18 | +1 |
| | A. thaliana X | 25 | 2769 | 2793 | 8/18 | +3 |
| ATG2 | yeast Atg2 | 18 | 1315 | 1332 | 6/18 | 0 |
| | human A | 18 | 1721 | 1738 | 6/18 | 0 |
| | human B | 18 | 1864 | 1881 | 6/18 | 0 |
| | A. thaliana | 18 | 1671 | 1688 | 7/18 | -4 |
| SHIP164 | human | / | | | | |
| | Dictyostelium | / | | | | |
| | A. thaliana | 41 | 1058 | 1098 | 7/18 | +1.5 |
| Tweek | yeast | 24 | 2933 | 2956 | 7/18 | +1 |
| | human | 40 | 4954 | 4993 | 7/18 | +1.5 |
| | Trichomonas | 24 | 2672 | 2695 | 8/18 | -1 |
| Hobbit | yeast Fmp27 | 20 | 2568 | 2586 | 8/18 | +3 |
| | yeast Hob2 | 20 | 2392 | 2411 | 5/18 | +1 |
| | human | / | | | | |
| | Fly | 22 | 2216 | 2237 | 5/18 | +2 |
| | A. thaliana | 18 | 2407 | 2424 | 6/18 | +5 |
