## Supplementary Data File 1 for "Sequence analysis and structural predictions of lipid transfer bridges in the repeating beta groove (RBG) superfamily reveals past and present domain variations affecting form, function and interactions of VPS13, ATG2, SHIP164, Hobbit and Tweek"

Supplementary Data File 1: Boundaries of RBG and other domains

Boundaries of all RBG domains and non-standard accessory domains in selected proteins from five RBG protein families. The C-terminal boundaries include the initial portion only (12-20 aa) of the 6$^{th}$ element (loop) which usually includes the start of the helix.

**VPS13 (22 proteins in 8 model organisms)**

human–A (3174 aa):– RBG domains (12): 1-145, 146-309, 480-609, 642-796, 877-1020, 1049-1181, 1205-1342, 1390-1527, 1549-1672, 1720-1855, 2518-2723 (omit WWE), 2762-2845; accessory domain: WWE (2553-2624)

human–B (4022 aa):– RBG domains (13): 1-149, 150-284, 386-612, 637-819, 838-966, 1064-1232, 1320-1519, 1546-1750, , 2116-2269, 2473-2612, 3380-3549, 3581-3639; accessory domain: β-sandwich (2295-2472)

human–C (3753 aa):– RBG domains (15): 1-191, 192-350, 524-658, 685-835, 906-1049, 1076-1214, 1236-1374, 1429-1558, 1589-1729, 1751-1889, 1940-2068, 2100-2223, 2274-2401, 3081-3282 (omit WWE), 3321-3402; accessory domain: WWE (3116-3181)

human–D (4388 aa):– RBG domains (15): 1-149, 150-300, 493-650, 704-904, 949-1130, 1158-1316, 1367-1556, 1611-1837, 1868-1990, 2017-2258, 2281-2418, 2461-2597, 2701-2814, 3820-3949, 3981-4050; accessory domains: UBA (2635-2676), Ricin (3569-3774)

fly–AC (3321 aa):– RBG domains (12): 1-149, 150-300, 476-620, 647-809, 899-1042, 1093-1224, 1245-1386, 1436-1581, 1623-1743, 1804-1934, 2663-2863, 2897-2977; accessory domain: WWE (2695-2769)

fly–B (3731 aa):– RBG domains (13): 1-149, 150-300, 338-536, 576-742, 759-865, 973-1120, 1182-1327, 1355-1534, 1685-1804, 1836-1943, 2168-2309, 3125-3278, 3316-3370; accessory domain: Ricin (2010-2167)

fly–D (3919 aa):– RBG domains (15): 1-149, 150-300, 475-612, 649-847, 874-986, 1007-1239, 1286-1422, 1456-1641, 1665-1789, 1814-1969, 2006-2140, 2164-2275, 2343-2471, 3357-3491, 3533-3592; accessory domains: UBA (2293-2338), Ricin (3120-3305)

worm–AC (3212 aa):– RBG domains (12): 1-147, 148-303, 488-594, 632-779, 842-966, 1038-1166, 1200-1332, 1381-1523, 1559-1679, 1748-1872, 2562-2765, 2800-2850; accessory domain: WWE (2594-2666)

worm–D (3312 aa):– RBG domains (11): 1-153, 154-301, 562-709, 733-976, 1001-1151, 1174-1344, 1373-1510, 1574-1716, 1745-1878, 2670-2857, 2888-2962; accessory domain: Ricin (2543-2647)

*Trichoplax*–AC (1561 aa):– RBG domains (/): 1-115, [RBG 2-9], 127-258, 899-1101, 1145-1222; accessory domain: WWE (936-1002)

*Trichoplax*–D (4149 aa):– RBG domains (15): 1-150, 151-302, 514-655, 694-890, 945-1096, 1119-1302, 1371-1520, 1546-1710, 1745-1863, 1891-2063, 2092-2226, 2273-

2410, 2509-2657, 3576-3624, 3739-3809; accessory domains: UBA (2430-2477), Ricin (3318-3513)

*S. cerevisiae* (3144 aa):– RBG domains (12): 1-157, 158-314, 505-645, 684-841, 890-1035, 1063-1184, 1210-1345, 1395-1525, 1560-1670, 1738-1855, 2579-2708, 2741-2819

*S. pombe–i* (3004 aa):– RBG domains (12): 1-149, 150-297, 462-587, 625-769, 811-949, 971-1090, 1116-1249, 1289-1416, 1450-1558, 1619-1736, 2438-2566, 2601-2676

*S. pombe–ii* (3071 aa):– RBG domains (12): 1-149, 150-306, 484-623, 662-815, 869-1008, 1036-1155, 1182-1315, 1352-1481, 1512-1623, 1678-1800, 2500-2631, 2662-2739

*Capsaspora*–AC (6160 aa):– RBG domains (13): 1-153, 154-401, 690-891, 929-1049, 1060-1420, 1440-2180, 2230-2370, 2450-2650, 2740-3060, 3110-3282, 3374-3496, 5030-5251, 5350-5507; accessory domains: helical (1199-2128), alpha/beta (unknown) (2845-3002)

*Capsaspora*–B (4668 aa):– RBG domains (13): 1-194, 195-400, 538-870, 902-1118, 1143-1312, 1380-1575, 1647-1870, 1937-2166, 2303-2460, 2518-2710, 2715-2890, 3940-4120, 4166-4222; accessory domains: helical (1675-1929), helical (2203-2298), helical (4327-4415)

*Capsaspora*–D (4768 aa):– RBG domains (15): 1-158, 159-307, 550-683, 719-1014, 1083-1246, 1311-1463, 1524-1654, 1736-1878, 1954-2119, 2148-2311, 2462-2602, 2726-2890, 3014-3138, 4164-4291, 4441-4514; accessory domains: helical (2936-3002), Ricin (3865-4090)

*Capsaspora*–X (3708 aa):– RBG domains (12): 1-156, 157-314, 540-670, 712-894, 972-1118, 1190-1319, 1376-1550, 1613-1746, 1823-1977, 2063-2191, 3124-3283, 3390-3538

*Arabidopsis*–S (3464 aa):– RBG domains (12): 1-149, 150-300, 464-598, 630-787, 802-952, 1055-1195, 1223-1376, 1429-1529, 1556-1731, 1748-1866, 2791-2941, 3017-3091

*Arabidopsis*–M1 (4219 aa):– RBG domains (12): 1-149, 150-300, 477-598, 633-780, 942-1159, 1197-1329, 1351-1488, 1533-1712, 1863-1974, 2395-2512, 3436-3565, 3584-3654; accessory domains: PH (801-909), beta-helix (1720-1860), beta-tripod (1983-2152), beta-tripod (2210-2374), C2 (2607-2765), beta-tripod (3972-4220)

*Arabidopsis*–M2 (4146 aa):– RBG domains (12): 1-169, 170-300, 533-657, 696-839, 1010-1238, 1281-1415, 1444-1573, 1615-1766, 1931-2038, 2440-2560, 3562-3702, 3711-3781; accessory domains: PH (874-989), beta-helix (1790-1930), beta-tripod (2039-2190), beta-tripod (2258-2420), C2 (2660-2820)

*Arabidopsis*–X (3125 aa):– RBG domains (9): 1-157, 158-314, 465-680, 730-900, 939-1181, 1207-1397, 1611-1742, 2558-2693, 2702-2776; accessory domains: Ricin (1428-1583), helices (partly amphiphathic) (2996-3125)

**ATG2**

human-B (2078 aa):- RBG domains (8): 1-132,163-335,505-701,738-930,1049-1218,1232-1363,1544-1726,1806-1869

*Arabidopsis* (1892 aa):- RBG domains (8): 1-120,172-325,457-639,661-869,895-1046,1062-1200,1428-1578,1612-1680

yeast (1592 aa):- RBG domains (8): 1-118,176-338,373-462,492-600,644-756,783-904,1025-1245,1262-1329

**SHIP164**

human (UHRF1BP1) (1464 aa):- RBG domains (6): 1-105,118-257,349-547,569-831,855-1300,1311-1370

*Arabidopsis* (1199 aa):- RBG domains (6): 1-126,127-290,315-466,481-654,688-843,902-1068

**Hobbit**

Human KIAA0100 (Hobbit, 2235 aa):- 1-134, 135-247, 276-393, 415-560, 568-685, 724-846, 869-1113, 1134-1293, 1318-1471, 1574-1774, 1907-2028, 2113-2166; accessory domain: paired helices 884-959, (1429-1444 – *i.e*. missing), 1781-1904

yeast Hob1 (Fmp27, 2628 aa):- RBG domains (12): 1-125, 125-225, 274-426, 467-593, 655-847, 893-1015, 1053-1403, 1429-1651, 1680-1985, 2070-2239, 2340-2470, 2522-2579; accessory domain: paired helices 1067-1234, 1835-1960, 2244-2339

**Tweek**

yeast Csf1:- RBG domains (17): 61-154, 226-448, 468-607, 627-768, 841-972, 993-1236 (omit loop with 2 helices 1081-1202), 1265-1389, 1534-1610, 1627-1745, 1778-1884, 1920-2035, 2077-2193, 2238-2320, 2354-2472, 2510-2671, 2712-2852, 2891-2922; accessory domains: helical bundle 1410-1520.
